# Structural mechanism of calcium-mediated hormone recognition and Gβ interaction by the human melanocortin-1 receptor

**DOI:** 10.1101/2021.06.20.449193

**Authors:** Shanshan Ma, Yan Chen, Antao Dai, Wanchao Yin, Jia Guo, Dehua Yang, Fulai Zhou, Yi Jiang, Ming-Wei Wang, H. Eric Xu

## Abstract

Melanocortins are peptide hormones critical for stress response, energy homeostasis, inflammation, and skin pigmentation. Their functions are mediated by five G protein-coupled receptors (MC1R to MC5R), predominately through the stimulatory G protein (Gs). MC1R, the founding member of melanocortin receptors, is mainly expressed in melanocytes and is involved in melanogenesis. Dysfunction of MC1R is associated with the development of melanoma and skin cancer. Here we present three cryo-electron microscopy structures of the MC1R-Gs complexes bound to endogenous hormone α-MSH, a marketed drug afamelanotide, and a synthetic agonist SHU9119. These structures reveal the orthosteric binding pocket for the conserved HFRW motif among melanocortins and the crucial role of calcium ion in ligand binding. They also demonstrate the basis of differential activities among different ligands. In addition, unexpected interactions between MC1R and the Gβ subunit were discovered from these structures. Together, our results provide a conserved mechanism of calcium-mediated ligand recognition, specific mode of G protein coupling, and a universal activation pathway of melanocortin receptors.

---

The melanocortin system is composed of five melanocortin receptors (MC1R to MC5R), four melanocortin-related peptide hormones, and two endogenous antagonists agouti and agouti-related peptide (AgRP)^1^. Melanocortins, with a highly conserved His-Phe-Arg-Trp (HFRW) sequence motif and consisting of adrenocorticotropic hormone (ACTH) and three melanocyte-stimulating hormones (α-MSH, β-MSH, and γ-MSH) (**Fig. 1a**), are derived from tissue-specific posttranslational processing of pro-opiomelanocortin (POMC)^2,3^. POMC is a precursor of polypeptide hormones, mainly secreted by the anterior pituitary, hypothalamus and brainstem^4^. The activity of POMC neurons is up-regulated by leptin and down-regulated by ghrelin, respectively^5^. Leptin, a satiety hormone, inhibits AgRP neurons and depolarizes POMC neurons to increase the expression of POMC and α-MSH. α-MSH, a 13-residue peptide hormone, was first identified in 1957 and is best known for maintaining energy homeostasis and protecting skin from ultraviolet radiation via augment of skin pigmentation^6–8^. Consequently, leptin decreases food intake and body weight by activating downstream signaling of the melanocortin system, while ghrelin, the hunger hormone, which is opposite to leptin, increases food intake and body weight by inhibiting melanocortin system signaling^9,10^. Dysregulation of melanocortins, leptin and ghrelin is associated with high risks of anorexia, cachexia and obesity^11–13^.

**Fig. 1.**
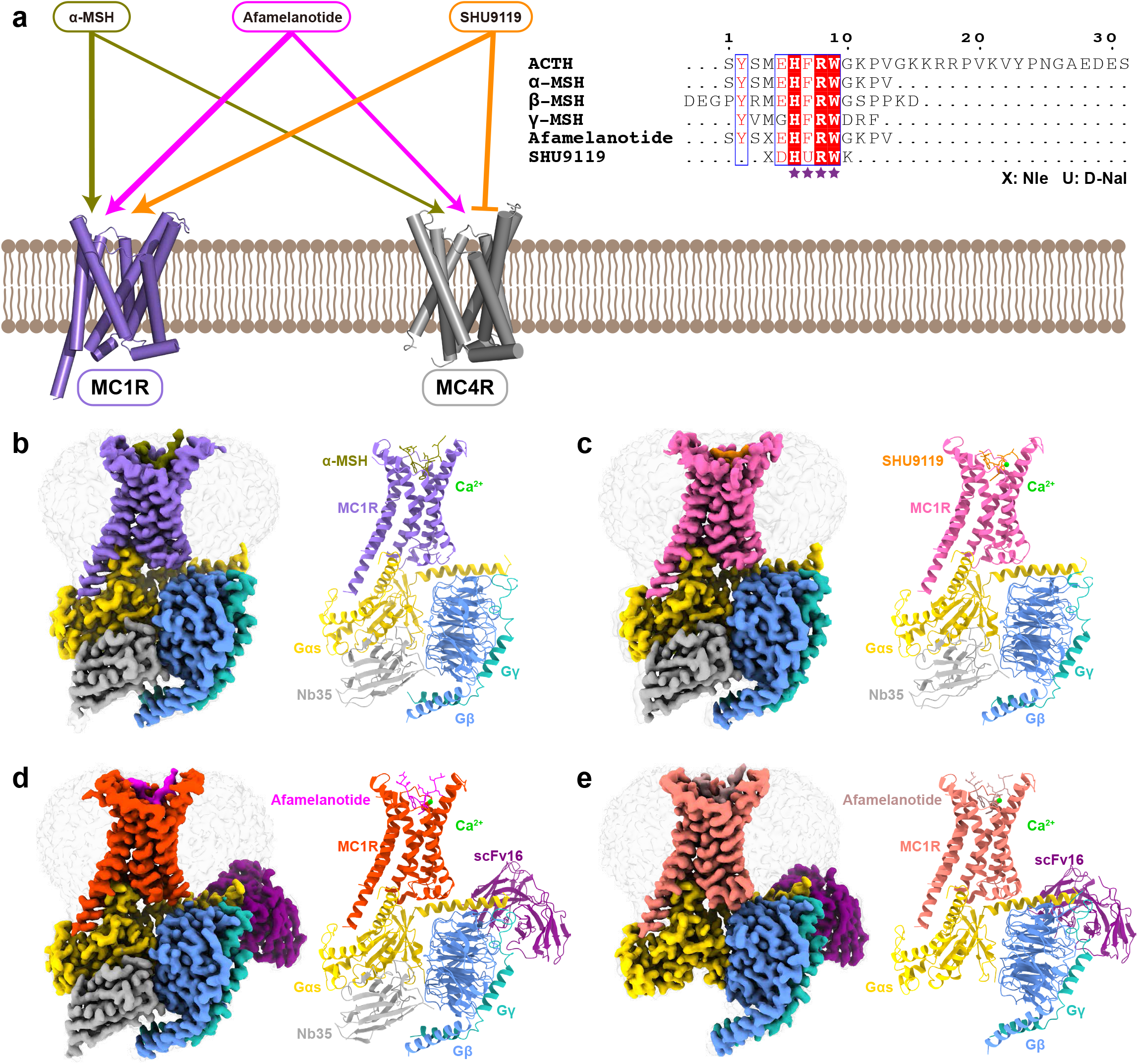
Cryo-EM structures of the MC1R-Gs complexes. **a** The left panel shows differential activities of α-MSH, afamelanotide and SHU9119 on MC1R and MC4R. The thickness of lines indicates the strength of affinity. The right panel is the sequence alignment of melanocortins with two synthetic peptides afamelanotide and SHU9119. The C-terminal residues of ACTH were omitted for clarify and the highly conserved HFRW motif is marked by purple stars. **b-e** Cryo-EM density maps (left panel) and cartoon representation (right panel) of the α-MSH-MC1R-Gs-Nb35 complex (**b**), SHU9119-MC1R-Gs-Nb35 complex (**c**), afamelanotide-MC1R-Gs-Nb35-scFv16 complex (**d**), and afamelanotide-MC1R-Gs-scFv16 complex (**e**). α-MSH is shown in olive, SHU9119 in dark orange, afamelanotide in magenta and rosy brown in both complexes. The corresponding MC1R is shown in medium purple, hot pink, orange red and salmon, respectively. Calcium ion is shown in lime, Gαs in gold, Gβ in cornflower blue, Gγ in light sea green, Nb35 in dark gray, and scFv16 in purple.

Activation of melanocortin receptors by cognate ligands induces a cascade of signal transduction through coupling to the stimulatory G protein (Gs) and arrestin^14^. MC1R to MC5R are among the shortest receptors in class A G protein-coupled receptors (GPCRs) that show distinct tissue-specific expression and physiological function. MC1R is mainly expressed in melanocytes and melanoma cells, and plays crucial roles in regulation of melanogenesis, skin pigmentation, and inflammation^15,16^. Abnormal functions of MC1R are linked to the development of melanoma and non-melanoma skin cancer^17–20^. MC2R is mostly located in the adrenal cortex and crucial for the hypothalamus-pituitary-adrenal (HPA) axis. Defective MC2R signaling causes a lethal disease called familial glucocorticoid deficiency (FGD)^21^. MC3R and MC4R, widely expressed in both the central nervous system and peripheral tissues, participate in the leptin-melanocortin signaling axis and are responsible for energy homeostasis, blood pressure, and inflammation. Selective ligands targeting MC3R and MC4R are promising drug candidates for obesity or anorexia^22–24^. MC5R is commonly seen in peripheral tissues and regulates exocrine gland secretion such as lacrimal, preputial and harderian glands^25^. However, structural basis for the complex interplay between melanocortins and MC1R-MC5R is largely unknown, except for the recent studies on MC4R^26–28^.

Given the important physiological functions of the melanocortin system, diverse synthetic ligands have been developed for therapeutic applications (**Fig. 1a**). Afamelanotide is the first synthetic α-MSH analog that has high affinity for MC1R^29^, and it has been approved as Scenesse™ by European Medicines Agency (EMA) for the prevention of phototoxicity in patients with erythropoietic protoporphyria^30^. SHU9119, a cyclic α-MSH analog, is a partial agonist for MC1R and MC5R but acts as an antagonist for MC3R and MC4R^31,32^. Currently, only the inactive crystal structure of MC4R bound to SHU9119 and the active cryo-electron microscopy (cryo-EM) structures of the MC4R-Gs complexes are available^26–28^. The limited structural information of the melanocortin system has hindered our understanding of the detailed mechanism by which various endogenous and synthetic peptides exert their differentiated actions. Here we present three cryo-EM structures of the MC1R-Gs complexes bound to α-MSH, afamelanotide and SHU9119 with a global resolution of 3.0 Å, 2.7 Å and 3.1 Å, respectively. The structures provide a paradigm for studying signal transduction of the melanocortin system and multiple structural templates for rational design of novel therapeutic agents targeting melanocortin receptors.

## RESULTS

### Cryo-EM structures of MC1R-Gs complexes

For cryo-EM studies, we co-expressed the full-length human MC1R, human dominant negative Gαs, human Gβ and human Gγ in High Five insect cells (**Supplementary information, Fig. S1a-b**). The structures of α-MSH, afamelanotide and SHU9119 bound MC1R-Gs complexes were determined at a resolution of 3.0 Å, 2.7 Å and 3.1 Å, respectively (**Supplementary information, Fig. S1c-e, S2 and Table S1**). In addition, a subset of afamelanotide bound MC1R-Gs complex without Nb35 were extracted and the structure was determined at a resolution of 2.9 Å (**Supplementary information, Fig. S1e and Table S1**). The high-quality EM maps allowed unambiguous model refinement of MC1R, the Gs heterotrimer and three bound peptide ligands α-MSH afamelanotide and SHU9119. Besides, a calcium ion was well defined in the EM maps (**Supplementary information, Fig. S3 and Table S1**).

Similarly, an annular detergent micelle surrounding the transmembrane domain (TMD) of MC1R was observed in all three structures mimicking the phospholipid bilayer. The receptors exhibit a nearly identical conformation with a large opening in the extracellular side of TMD (**Fig. 1b-e**). Different from other class A GPCRs, the extracellular loop 2 (ECL2) of MC1R is extremely short and its ECL3 forms an ordered helix (**Fig. 2**). Three peptides adopt a U shape conformation in the extracellular end of the TMD with a similar orientation. In addition, the well-defined calcium ion near TM3 is positioned to stabilize MC1R ligand binding.

**Fig. 2.**
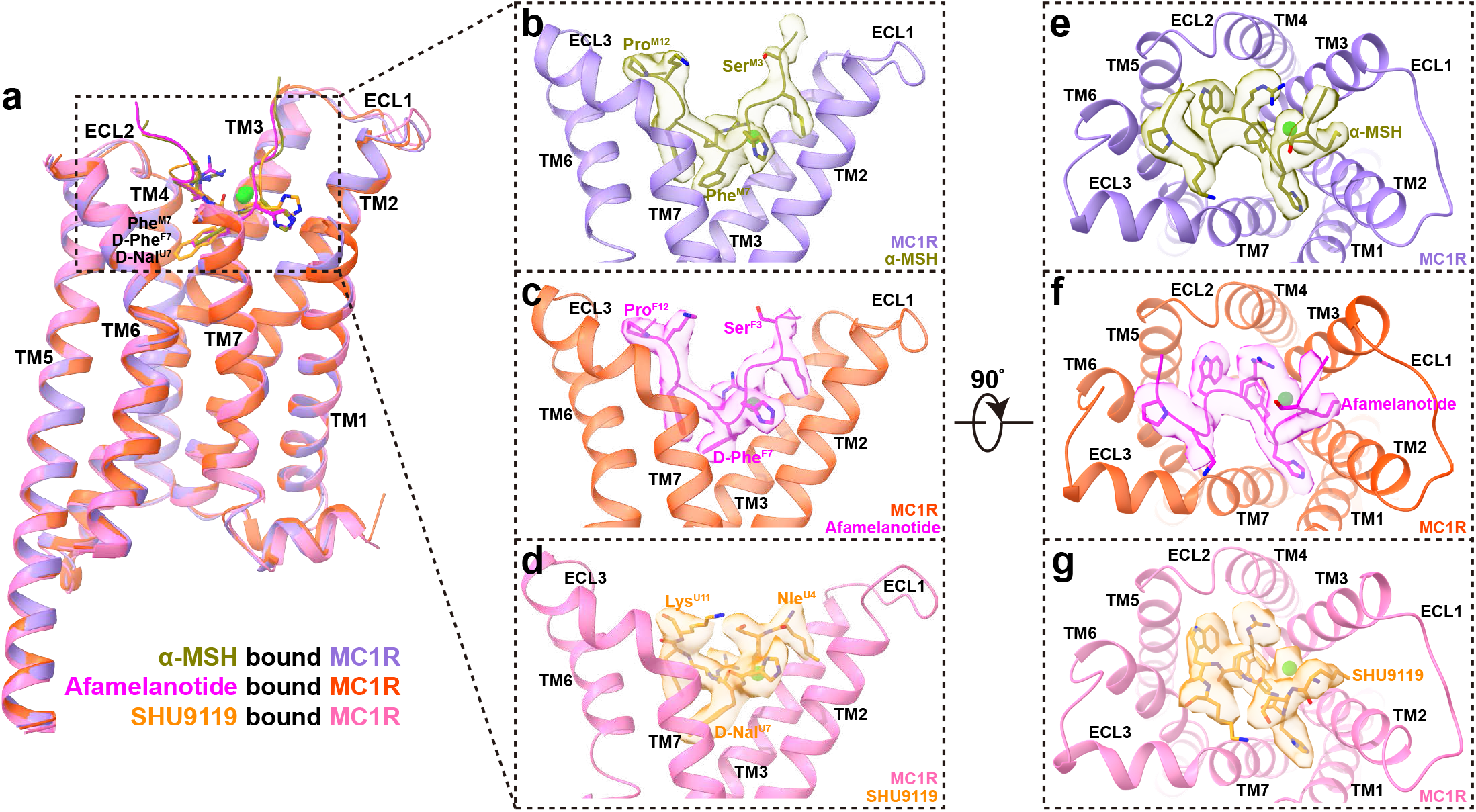
The orthosteric peptide binding pocket of MC1R. **a** Structural comparison of α-MSH bound MC1R, afamelanotide bound MC1R and SHU9119 bound MC1R. ECD and ECL3 of MC1R were omitted for clarify and the alignment was based on the receptor. **b-g** Side views (**b**-**d**) and top views (**e**-**g**) of the orthosteric binding pocket in α-MSH bound MC1R complex (**b**, **e**), afamelanotide bound MC1R complex (**c**, **f**) and SHU9119 bound MC1R complex (**d**, **g**). Calcium ion is displayed in sphere and colored in lime. The EM density maps of α-MSH, afamelanotide, SHU9119 and calcium ion are shown at 0.08 threshold.

### Orthosteric peptide binding pocket

The overall structures of the three MC1R-Gs complexes are highly similar with root mean square deviation (RMSD) values of 0.70 Å for the Cα atoms between α-MSH and afamelanotide bound MC1R, and 0.87 Å for the Cα atoms between α-MSH an SHU9119 bound MC1R (**Fig. 2a**). All three peptides adopt a U shape conformation in the extracellular end of TMD, with the benzene ring of Phe^M7/F7^ and the naphthalene ring of D-Nal^U7^ penetrating deeply into the TMD core (superscript M refers to α-MSH, F to afamelanotide and U to SHU9119, residue numbers are based on α-MSH) (**Fig. 2b-d**). The interactions of α-MSH, afamelanotide and SHU9119 with MC1R bury a total interface area of 2085 Å^2^, 1986 Å^2^ and 1790 Å^2^, respectively (**Fig. 2e-g**). The smaller interface area between SHU9119 and MC1R might explain why SHU919 is a weaker agonist than α-MSH and afamelanotide for MC1R (**Supplementary information, Fig. S4a and Table S5**). The highly conserved HFRW motif of melanocortins is at the center of the U shape pocket and provides the major contacts for binding to MC1R (**Fig. 3**).

**Fig. 3.**
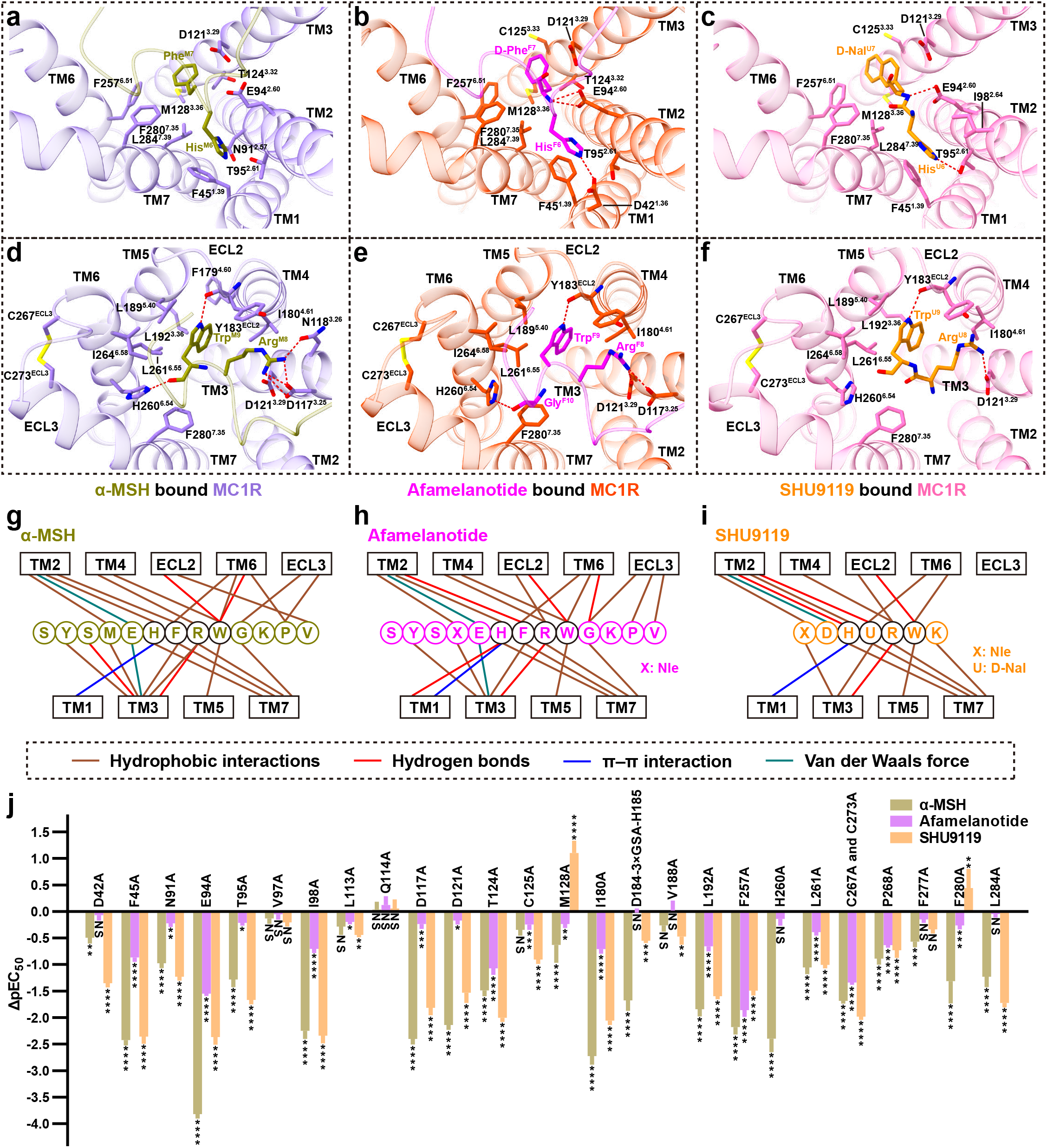
Molecular recognition of α-MSH, afamelanotide and SHU9119 by MC1R. **a-f** Two different views of the detailed interactions between α-MSH and MC1R (**a**, **d**), afamelanotide and MC1R (**b**, **e**), SHU9119 and MC1R (**c**, **f**). **a**-**c** show interactions of His^6^ and Phe^M7^/D-Phe^F7^/D-Nal^U7^ in peptide ligands with MC1R. **d**-**f** depict interactions of Arg^8^ and Trp^9^ in peptide ligands with MC1R. C267^ECL3^ of MC1R forms a conserved disulfide bond with C273^ECL3^. Hydrogen bonds are shown as red dash lines. **g-i** Schematic diagrams of the interactions between α-MSH and MC1R (**g**), afamelanotide and MC1R (**h**), SHU9119 and MC1R (**i**). The highly conserved residues in peptides are surrounded by black circles. **j** Effects of mutations in the orthosteric binding pocket of MC1R on α-MSH, afamelanotide and SHU9119 induced cAMP accumulation. Values are presented as means ± S.E.M. from at least three independent experiments performed in quadruplicate. All data were analyzed by one-way ANOVA and Dunnett’s test. *P<0.05, **P<0.01, ***P<0.001 and ****P<0.0001. NS, not significant (comparison between the wild-type (WT) MC1R and its mutants).

The orthosteric peptide-binding pocket can be divided into three parts based on the conformation of α-MSH (**Fig. 2**). The first part is formed by the N-terminal residues 1-5, which is parallel to the plane between TM2 and TM3. The second part is the critical HFRW motif (residues 6-9), which is inserted deeply into the TMD core and interacts with TM1 to TM7. The third part is formed by the C-terminal residues 10-13, which is in proximity to TM6, TM7, and ECL3 (**Fig. 2**). Extensive hydrophobic and polar interactions are observed between MC1R and three peptides (**Fig. 3 and supplementary information, Table S2-S4**) and the majority of residues involved in peptide binding are conserved in melanocortin receptors. For example, His^6^ of peptides packs against F45^1.39^ (superscripts denote the Ballesteros-Weinstein numbers^33^) forming a conserved π-π interaction (**Fig. 3a-c**). The positively charged side chain of Arg^8^ of peptides forms hydrogen bonds with the negatively charged side chains of D117^3.25^ or D121^3.29^ (**Fig. 3d-f**). In order to correlate these structural observations with signaling profiles, various mutants were constructed to detect cAMP responses of MC1R (**Fig. 3j and Supplementary information, Fig. S4b-d, Table S5-S6**). The majority of alanine mutations in the orthosteric peptide-binding pocket reduced both potency and efficacy of ligand-stimulated cAMP accumulation mediated by MC1R. Notably, there is a considerable divergence in the basal activities of different MC1R constructs, consistent with the constitutive activity of MC1R reported previously^34–36^. Therefore, the decline in pEC_50_ values and cAMP responses of MC1R mutants elicited by three peptides reveal an important role of these residues in ligand binding and receptor activation (**Fig. 3j and Supplementary information, Fig. S7**).

Of note is the observation of an extremely short ECL2 in the three MC1R-Gs complexes, different from a longer ECL2 in the β_2_AR-Gs complex, where it forms a lid covering the extracellular top of TMD (**Supplementary information, Fig. S4e**)^37^. In the case of GPR52, ECL2 can fold into the transmembrane bundle and function as a built-in ‘agonist’ to activate the receptor^38^. In addition, TM2 and ECL1 of MC1R move outwards compared to β_2_AR. In the MC1R-Gs complex structures, ECL3 forms an ordered helix, in which two cysteines, C267^ECL3^ and C273^ECL3^, make a disulfide bond instead of the canonical disulfide bond between TM3 and ECL2 seen in other class A GPCRs (**Fig. 3d-f**). Such a unique feature of the MC1R structure allows a broader opening in the extracellular side of TMD to accommodate larger peptide ligands and a calcium ion. Extending ECL2 by a nine-residue insertion (3×GSA) between D184^ECL2^ and H185^ECL2^ or disruption of the ECL3 disulfide bond through mutations of C267^ECL3^A and C273^ECL3^A decreased both pEC_50_ values and potencies of the three peptides (**Supplementary information, Fig. S4f-g and Table S5-S6**).

### Differential activities of peptide ligands

The three peptide ligands (α-MSH, afamelanotide and SHU9119) used in this study display differential activities toward different melanocortin receptors, which can be readily explained by our structural observations. Specifically, afamelanotide, which has D-Phe^F7^ instead of Phe^M7^ in α-MSH, has a higher affinity for melanocortin receptors. SHU9119 with D-Nal^U7^ displays a partial agonism for MC1R and MC5R but acts as an antagonist for MC3R and MC4R. Structural analysis of the ligand-binding pocket of MC1R reveals that the change of Phe^M7^ causes slightly different orientations of nearby residues, resulting in different interactions between MC1R and the peptides (**Fig. 3 and Fig. 4a-b**). For example, the hydroxyl group on the carboxyl group of Trp^M9^ forms a hydrogen bond with the nitrogen on the imidazole ring of H260^6.54^, which is absent in the afamelanotide bound MC1R-Gs complex (**Fig. 3d-e**). H260^6.54^A mutation decreases the affinity of α-MSH for MC1R, without affecting that of afamelanotide (**Supplementary information, Fig. S4d, g and Table S5-S6**). Besides, in comparison with the inactive SHU9119-MC4R complex, the benzene ring of Phe^M7^ in the active α-MSH-MC1R-Gs complex inserts into the TMD and induces a downward shift of F257^6.51^ and F280^7.35^ of MC1R, which makes steric clash with the toggled switch residue W254^6.48^ and pushes W254^6.48^ into the active position (**Fig. 4a**). The rearrangement of the toggled switch residue W254^6.48^ is a molecular hallmark to start a cascade of conformational changes during receptor activation. However, the cyclic structure of SHU9119, which is different from non-cyclic peptides of α-MSH and afamelanotide, makes more compact interactions with MC1R and a smaller shift of F257^6.51^ and F280^7.35^ (**Fig. 4b**), providing a basis for the partial agonism of SHU9119 toward MC1R, in which α-MSH and afamelanotide are full agonists.

**Fig. 4.**
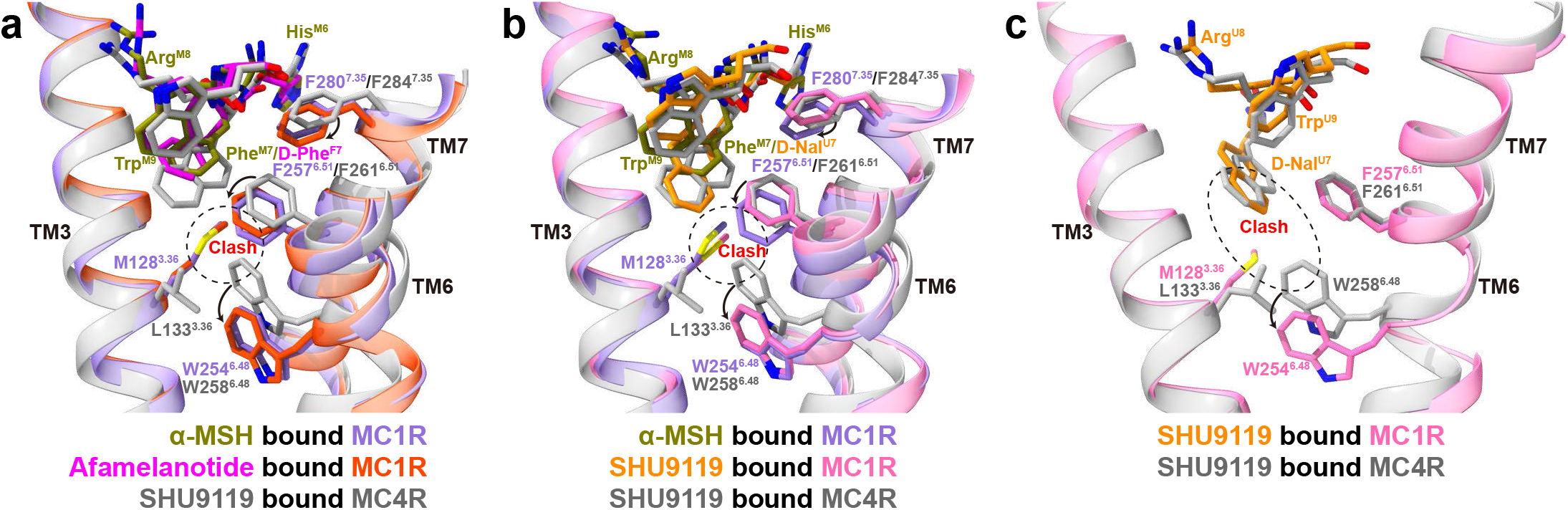
Structural basis of differential activities among melanocortin peptide ligands. **a-b** Structural comparison of α-MSH bound MC1R complex (**a**-**b**), afamelanotide bound MC1R complex (**a**), SHU9119 bound MC1R complex (**b**) and SHU9119 bound MC4R complex (PDB: 6W25, **a**-**b**). The conformational changes of W254^6.48^, F257^6.51^ and F280^7.35^ upon MC1R activation are emphasized. The alignment was based on the receptors and MC1R residue numbers are colored in medium purple. **c** Comparison of SHU9119-binding pocket in MC1R and MC4R. M128^3.36^ of MC1R forms steric clash with D-Nal^U7^ and W254^6.48^, making SHU9119 a partial agonist for MC1R. Hydrogen bonds are shown as red dash lines.

In addition, when comparing the structures of SHU9119 bound MC1R with SHU9119 bound MC4R, several differences are observed in their ligand-binding pockets despite that most pocket residues are conserved. Notably, D-Nal^U7^ of SHU9119 was constrained in the TMD core by L133^3.36^ and F261^6.51^ of MC4R in the inactive SHU9119-MC4R complex structure^26^. However, L133^3.36^ of MC4R corresponds to M128^3.36^ in MC1R. The side chain of M128^3.36^ moves upward to interact with D-Nal^U7^, causing a severe steric clash with the toggled switch residue W254^6.48^ and a subsequently downward movement of W254^6.48^ (**Fig. 4c**). M128^3.36^L mutation of compromised SHU9119-stimulated cAMP response of MC1R (**Supplementary information, Fig. S5a and Table S5**), consistent with that L133^3.36^M mutation of MC4R converted SHU9119 from an antagonist to a partial agonist^27,39^. Together, these results provide a basis of SHU9119 as an agonist for MC1R and as an antagonist for MC4R.

### Role of calcium ion

Extensive evidence reveal that the divalent ion is of crucial importance for melanocortin signaling. Calcium ion assists melanocortins in binding to their cognate receptors with a better effect than magnesium ion^40–42^. Zinc ion activates MC1R and MC4R by acting as an agonist or allosteric modulator^43,44^. A well-resolved electron density of calcium ion was observed in the MC1R-Gs complexes at the same position as that of the SHU9119 bound MC4R structure (**Fig. 5a-c**). The Ca^2+^-binding pocket is conserved within the orthosteric peptide binding pocket, consisting of E^2.60^, D^3.25^, and D^3.29^ from melanocortin receptors (**Fig. 5d**) as well as Glu/Asp^5^, Phe^7^ and Arg^8^ from melanocortins. Declined cAMP responses and peptide affinities for MC1R with mutations of E94^2.60^A, D117^3.25^A and D121^3.29^A are likely the consequence of destroying both peptide and calcium ion binding pockets (**Supplementary information Fig. S5c-d and Table S5-S6**). The affinity of α-MSH for MC1R increases when Ca^2+^ concentrations are elevated (**Supplementary information Fig. S5b**). Specifically, addition of 0.5 mM Ca^2+^ shifted cAMP response curve to the left upon stimulation with α-MSH and SHU9119 (500-fold) or afamelanotide (10-fold) (**Supplementary information Fig. S5e**), pointing to an allosteric modulation role of Ca^2+^.

**Fig. 5.**
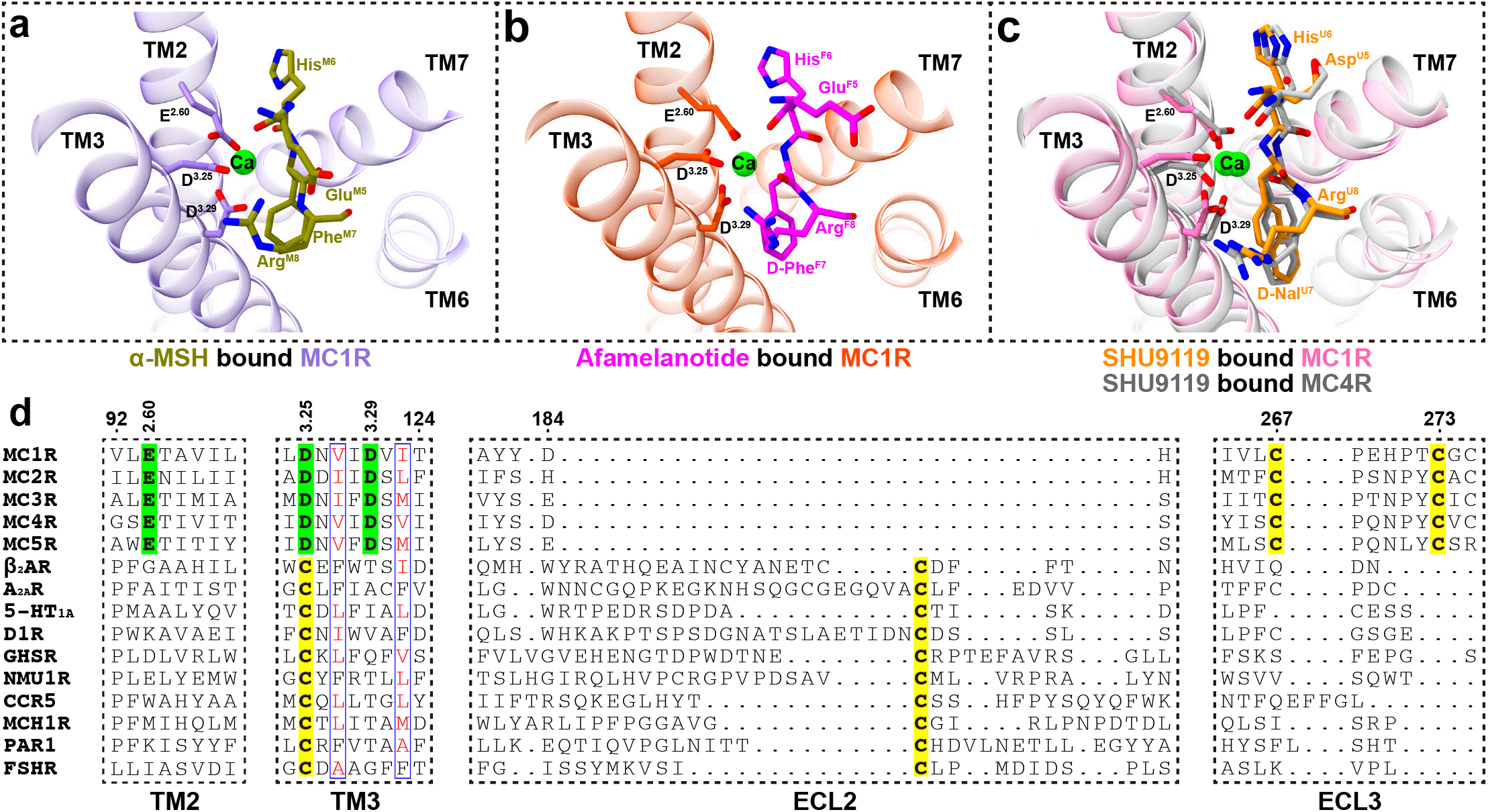
The calcium binding pocket of melanocortin receptors. **a-c** Expanded views of the calcium ion binding pocket in α-MSH bound MC1R complex (**a**), afamelanotide bound MC1R complex (**b**), SHU9119 bound MC1R and MC4R complexes (**c**). The alignment was based on the structures of MC1R and MC4R. **d** Sequence alignment of melanocortin receptors with other class A GPCRs from different branches of the rhodopsin family. The residues involved in the calcium binding pocket are highlighted in green and the cysteines forming conserved disulfide bonds are highlighted in yellow.

It is noteworthy that D^3.25^ of melanocortin receptors corresponds to highly conserved C^3.25^ which forms a canonical disulfide bond with cysteine of ECL2 in other class A GPCRs (**Fig. 5d**). However, the extremely short ECL2 and the calcium-binding pocket of MC1R preclude the possibility of a disulfide bond between ECL2 and TM3. Instead, two cysteines of ECL3 form a conserved disulfide bond in melanocortin receptors, which was absent in other class A GPCRs (**Supplementary information, Fig.S5f**). These distinct features demonstrate that the calcium-binding pocket is both conserved and unique in all five melanocortin receptors.

### Activation of MC1R

The active SHU9119-MC1R-Gs complex reported here together with that of previous inactive structure of SHU9119-MC4R complex reveal large conformational changes upon receptor activation (**Fig. 6a**). At the extracellular side, ligand binding induced an inward movement of TM1 by 1.7 Å at F45^1.39^ and an outward movement of TM2 by 2.2 Å at L101^2.67^ (**Fig. 6b**). At the cytoplasmic side, TM3, TM4 and TM7 moved inwards slightly, TM5 extended by four helices and moved inwards to interact with Gs, while TM6 moved outwards by 13.4 Å at L237^6.31^ (**Fig. 6c**). The pronounced outward movement of TM6 in MC1R is consistent with that seen among activated Gs coupled receptors.

**Fig. 6.**
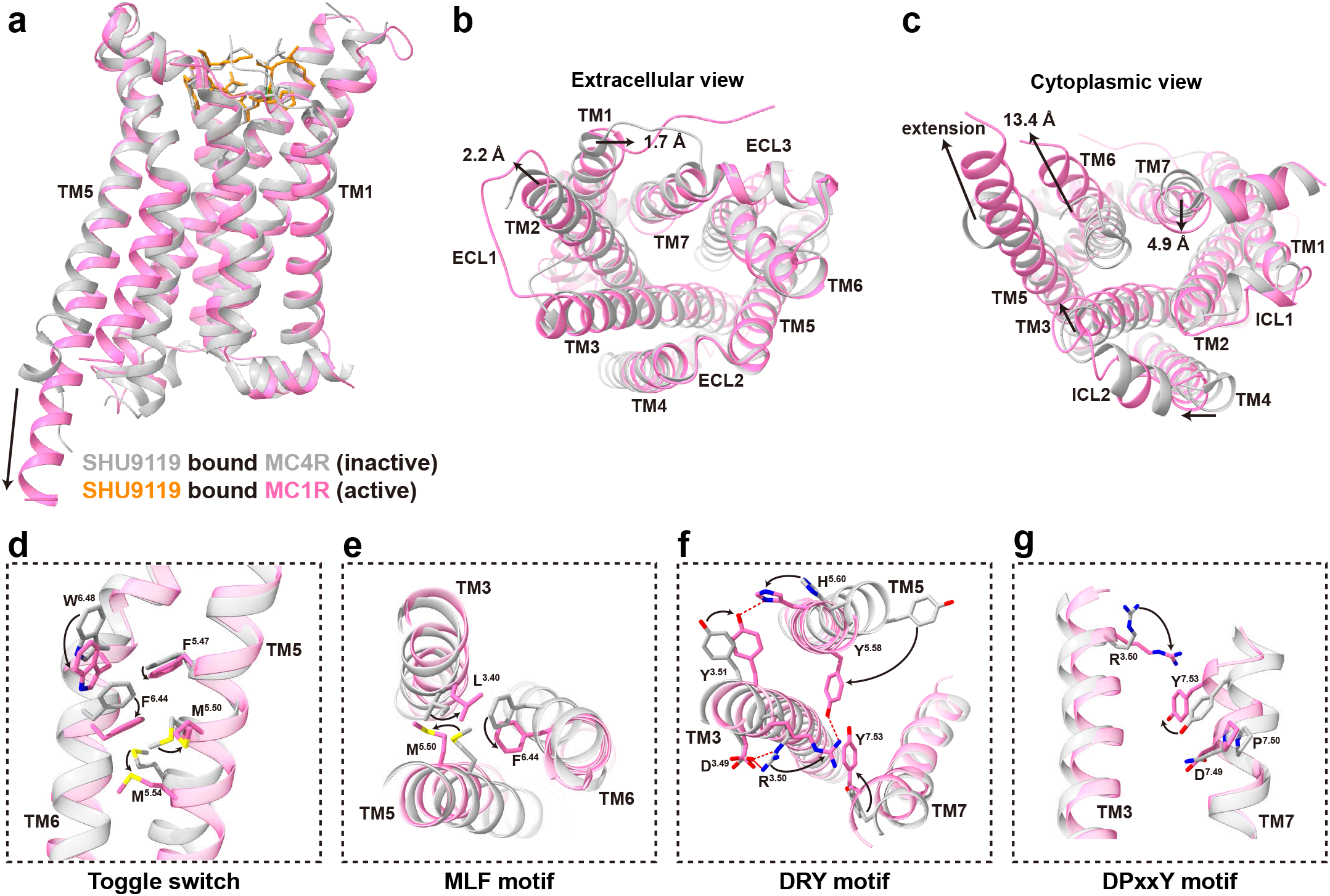
Activation of MC1R by peptide ligands. **a-c** Structural alignment of the SHU9119 bound MC1R and MC4R complexes (PDB: 6W25). The alignment was based on the structures of MC1R and MC4R, which are colored hot pink and dark gray, respectively. **a** side view; **b** extracellular view; **c** intracellular view. **d-g** Conformational changes of the conserved “micro-switches” upon receptor activation. **d** Toggle switch; **e** MLF motif; **f** DRY motif; **g** DPxxY motif. The conformational changes of residue side chains are shown as arrows. Hydrogen bonds are shown as red dash lines.

As mentioned above, the Phe^7^ of melanocortin peptide ligands interacts with M128^3.36^, inducing a downward movement of the toggled switch residue W254^6.48^ and a subsequently downward movement of F250^6.44^ (**Fig. 6d-e**). In contrast to the conserved P^5.50^I^3.40^F^6.44^ motif, M199^5.50^ in melanocortin receptors fits better in α-helical conformation than P^5.50^, generating a straight helix without the bulge as observed in the β2AR-Gs complex (**Fig. 6e and supplementary information, Fig. S6a**). Structural superimposition with the inactive structure of SHU9119-MC4R complex reveals that M199^5.50^ in the active MC1R-Gs complex changes its orientation to induce an inward movement of TM5.

In addition, the highly conserved D^3.49^R^3.50^Y^3.51^ motif in class A GPCRs is shown to be critical for receptor activation (**Fig. 6f**). Upon activation, Y143^3.51^ moves inwards to form hydrophobic interactions with TM5 and a hydrogen bond with H209^5.60^. The side chain of R142^3.50^ stretches out straightly, breaking the salt bridge with D141^3.49^ and pushing TM6 away from the TMD core. Meanwhile, R142^3.50^ packs against Tyr391 of Gαs (**Fig. 7a**) and contributes to stable interactions with Y207^5.58^ and Y298^7.53^. The DRY motif links the cytoplasmic ends of TM3, TM5, TM7 and Gαs, playing a direct role in stabilizing the active state of MC1R.

**Fig. 7.**
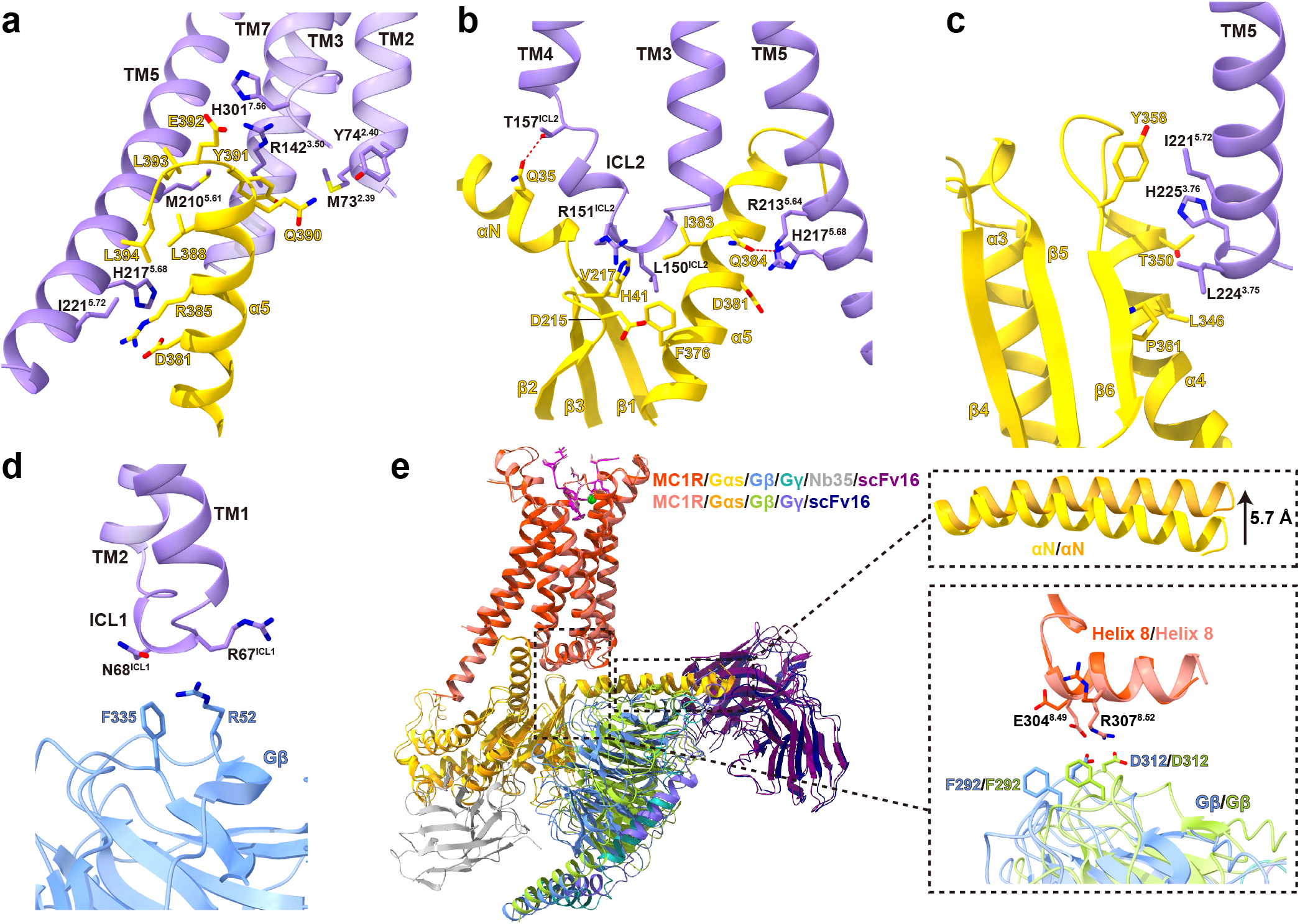
G protein coupling of MC1R. **a** The interactions between MC1R (medium purple) and α5 helix of Gαs (gold) in the cavity at the cytoplasmic region of MC1R. **b** The interactions between ICL2 of MC1R and Gαs. ICL2 inserts into the groove formed by αN-β1 hinge, β2-β3 loop and α5 helix of Gαs. **c** The interactions between the C terminus of TM5 and α4 helix, α4-β6 loop and β6 sheet of Gαs. **d** The interactions between ICL1 of MC1R and Gβ. **e** Comparison of Gs protein coupling between afamelanotide-MC1R-Gs-Nb35-scFv16 complex and afamelanotide-MC1R-Gs-scFv16 complex. The alignment was based on the receptor. Differences are in the αN helix of Gαs and the interactions between helix 8 of MC1R and Gβ.

Melanocortin receptors contain an aspartate (D^7.49^) instead of an asparagine (N^7.49^) at the conserved N^7.49^P^7.50^xxY^7.53^ motif as seen in most class A GPCRs (**Fig. 6g**). D294^7.49^N mutation exhibited a nearly equivalent cAMP response of MC1R stimulated by α-MSH, while D294^7.49^A mutation remarkably impaired the peptide’s ability to activate MC1R, suggesting that both DPxxY and NPxxY motifs could effectively govern the transition of GPCRs from inactive to active states (**Supplementary information, Fig. S6e and Table S5**). Taken together, MC1R activation involves a cascade of conformational changes through rearrangement of the toggle switch W^6.48^, P(M)I(L)F, DRY and N(D)PxxY motifs present in most class A GPCRs.

### Unique features of G protein coupling by MC1R

The massive conformational changes in the cytoplasmic side of TMD is supposed to accommodate α5 helix of Gαs, which is the primary structural element interacting with MC1R. There is negligible difference in Gs coupling among α-MSH, afamelanotide and SHU9119 bound MC1R, and the interactions of Gs with MC1R bury a total surface area of 3252 Å^2^, 3166 Å^2^ and 3044 Å^2^, respectively. α5 helix of Gαs inserts into the cytosolic core surrounded by TM3, TM5, TM6, TM7 and ICL2 (**Fig. 7a**). R142^3.50^ packs against Tyr391 of Gαs, stabilized by van der Waals interaction. H301^7.56^ forms a salt bridge with Glu392 of Gαs and mutating H301^7.56^ to alanine impairs the basal cAMP activity of MC1R (**Supplementary information, Fig. S6f and Table S5**). The extension of TM5 allows further interactions with Gαs (**Fig. 7a-c**). M210^5.61^, R213^5.64^, H217^5.68^ and I221^5.72^ of TM5 make substantial polar and hydrophobic interactions with α5 helix of Gαs. The C terminus of TM5 directly contacts α4 helix, α4-β6 loop and β6 sheet of Gαs (**Fig. 7c**). Alanine mutations of the C-terminal residues of TM5 result in a significant reduction of cAMP responses (**Supplementary information, Fig. S6f and Table S5**).

Furthermore, the intracellular loops facilitate additional interactions with Gs to stabilize the complex. Typically, ICL2 adopts a 3^10^-helix conformation and inserts into the groove formed by αN-β1 hinge, β2-β3 loop and α5 helix of Gαs (**Fig. 7b**). It was reported that the binding of ICL2 to Gαs induces sequential activation of Gs to release GDP^45^. The side chain of L150^ICL2^ is enclosed by the hydrophobic interactions with His41, Val217, Phe376 and Ile383 of Gαs. L150^ICL2^A mutant dramatically suppressed the ability of MC1R to couple Gs to elicit cAMP response (**Supplementary information, Fig. S6f and Table S5**). Different from class B GPCRs, ICL1, rather than helix 8, forms van der Waals interactions with Arg52 and Phe335 of Gβ (**Fig. 7d**)^46^. Mutating residues of ICL1 to alanine destabilizes the complex and impairs the cAMP response of MC1R (**Supplementary information, Fig. S6g and Table S5**).

Interestingly, compared to the afamelanotide bound MC1R-Gs-Nb35-scFv16 complex (the Nb35^plus^ complex), Gs in the absence of Nb35 (the Nb35^minus^ complex) adopts a relatively loose conformation as Nb35 interacts with both Gαs and Gβ (**Fig. 7e**). The αN helix of Gαs moved upwards by 5.7 Å despite the nearly overlapping α5 helix. The most surprising observation is that Gβ from the Nb35^minus^ complex is in proximity to MC1R to make direct contacts with E304^8.49^ and R307^8.52^ of helix 8 (**Fig. 7e and supplementary information, Fig. S6b-c**). Deletion of helix 8 deprived the ability of the three peptides to activate MC1R and dual mutations of E304^8.49^A and R307^8.52^A considerably weakened the Gs coupling, which was consistent with that the mutations in the corresponding residues of Gβ decreased the efficacy of G protein activation induced by three peptides α-MSH, afamelanotide and SHU9119 (**Supplementary information, Fig. S6h-i and Table S5**). Such a rearrangement results in a higher interface area of 3764 Å^2^ between MC1R and Gs in the Nb35^minus^ complex. No obvious shift was observed on peptide binding and receptor activation between the Nb35^plus^ and the Nb35^minus^ complexes.

## DISCUSSION

In this paper, we present three active cryo-EM structures of the MC1R-Gs complexes bound to α-MSH, afamelanotide or SHU9119. These structures reveal a unique orthosteric peptide-binding pocket in the extracellular side of MC1R, where a wide opening is observed and allowed by the extremely short ECL2 and the ordered ECL3 to hold relatively large peptide-hormones. In this pocket, three peptides adopt a similar U-shape conformation, with the highly conserved HFRW motif among melanocortins lying at the bottom. It is noteworthy that this motif makes major contacts with MC1R and provides indispensable energy to stabilize the binding with MC1R. The binding mode between three peptides and MC1R is similar to that of recently reported setmelanotide-MC4R-Gs complex (**Supplementary information, Fig. S6j**), indicating a conserved mechanism of ligand recognition by different family members of melanocortin receptors^27^.

Structural superimposition of the three MC1R-Gs complexes shows that the critical residue Phe in the HFRW motif contributes to the differential activities among three melanocortin peptides. Substitution of Phe^M7^ with D-Phe^F7^ or D-Nal^U7^ affects the orientation of nearby residues and the extent of peptide insertion into the binding pocket, thereby distinguishing the detailed interactions between different peptides and MC1R. Accordingly, mutations in this binding pocket led to different effects of the peptides used with afamelanotide being the strongest agonist as it was least affected. Particularly, we demonstrate that M128^3.36^ of MC1R, instead of L133^3.36^ of MC4R, is a key residue that converts SHU9119 from an antagonist of MC4R to a partial agonist of MC1R.

Notably, Ca^2+^ was observed in all three MC1R-Gs complexes. Sequence alignment and structural comparison among melanocortin receptors and with other class A GPCRs highlight a unique and conserved calcium-binding pocket consisting of E^2.64^, D^3.25^ and D^3.29^ of melanocortin receptors and the backbone of melanocortin peptides. Specifically, the existence of Ca^2+^ excludes the canonical disulfide bond between TM3 and ECL2 seen in other class A GPCRs. Depletion of Ca^2+^ or disruption of the calcium-binding pocket reduced the potencies and efficacies of cAMP responses elicited by three peptides and mediated by MC1R, indicating a crucial role of the calcium ion in ligand recognition and MC1R activation.

It is known that the residues involved in the receptor activation and Gs coupling are conserved, delineating a universal mechanism among class A GPCRs. However, except for the interactions between ICL1 and Gβ as seen in other GPCR-Gs complexes^38,47,48^, unexpected interactions between helix 8 of MC1R and Gβ were found in the Nb35^minus^ complex. Structural superimposition of the Nb35^minus^ complex with the Nb35^plus^ complex reveals that the Gs heterotrimer protein adopts a relatively loose conformation and Gβ is closer to the receptor to make direct interactions with helix 8 of MC1R in the absence of Nb35. Mutations in helix 8 and Gβ both markedly reduced the potency of Gs coupling by MC1R, suggesting that the interaction between helix 8 and Gβ is important for G protein coupling. To date, such interactions have only been observed in D1R and some class B GPCRs^46–57^. The residues from helix 8 of MC1R that form interactions with the Gβ subunit are conserved in D1R, implying that Gβ interaction may be a shared feature of G protein recruitment by certain GPCRs (**Supplementary information, Fig. S6b-d**). Since most class A GPCR-Gs complex structures were solved with Nb35 and the interaction between the Gβ subunit and helix 8 in MC1R was only observed in the Nb35^minus^ complex, it is suggested that other class A GPCRs could also interact with the Gβ subunit through their helix 8.

## MATERIALS AND METHODS

### Constructs of MC1R and Gs

The full-length human MC1R was cloned into pFastBac vector (Invitrogen) with its native signal peptide replaced by the haemagglutinin (HA) signal peptide followed by a 10 × His tag and cytochrome b562RIL (BRIL) as a fusion partner. To facilitate expression and purification, the C terminus of MC1R was fused with a 15-amino-acid polypeptide linker (GSSGGGGSGGGGSSG) and a LgBiT (Promega).

Human Gαs was constructed based on miniGs (PDB: 5G53) deleting switch III and including eight mutations (G49D, E50N, L63Y, A249D, S252D, L272D, I372A and V375I)^58^. Two additional dominant-negative mutations (G226A and A366S) were introduced to Gαs to decrease the affinity of nucleotide binding and increase the stability of the heterotrimeric G protein^59^. The N terminus (M1-K25) and α-helical domain (AHD, G67-L203) of Gαs were replaced by the N terminus (M1-M18) and AHD (G60-K180) of human Gαi, which was initially designed to bind scFv16 and Fab_G50^60,61^. Human Gβ with a C-terminal 15-amino-acid polypeptide linker followed by a HiBiT (peptide 86, Promega) and human Gγ were cloned into pFastBac vector, respectively ^62^. scFv16 was constructed into the same vector with an N-terminal GP67 signaling peptide.

### Preparation of Nb35

Nb35 was expressed and purified according to previously described methods^63^. The purified Nb35 was concentrated and stored in −80°C.

### Expression and purification of the MC1R-Gs complex

Recombinant viruses of MC1R, Gαs, Gβ, Gγ and scFv16 were generated using Bac-to-Bac baculovirus expression system (Invitrogen) in *sf*9 insect cells (Expression Systems). High Five™ cells (ThermoFisher) at a density of 2 × 10^6^ cells/mL were transfected with above five baculoviruses at a ratio of 1:1:1:1:1. The cells were cultured for 48 h at 27°C after infection and collected by centrifugation at 2 × 1000 rpm for 20 min. Notably, the α-MSH-MC1R-Gs and SHU9119-MC1R-Gs complexes were expressed without scFv16.

The cell pellets were suspended in 20 mM HEPES, 100 mM NaCl, 100 μM TCEP, pH 7.4, supplemented with protease inhibitor cocktail (EDTA-Free) (Bimake) and centrifuged at 30,000 × *g* for 30 min. The pellets were lysed in the same buffer supplemented with 40 mM imidazole, 10 mM MgCl_2_ and 5 mM CaCl_2_, and the complex formation was initiated by addition of 25 mU/mL Apyrase (Sigma), 20 mg/mL Nb35 and 10 μM peptide (GenScript). The lysate was incubated for 1.5 h at room temperature (RT) followed by addition of 0.5% (w/v) lauryl maltose neopentylglycol (LMNG, Anatrace) and 0.1% (w/v) cholesterol hemisucinate (CHS, Anatrace) for 3 h at 4°C to solubilize the membrane. The supernatant was isolated by centrifugation at 65,000 × *g* for 30 min and incubated with Ni-NTA beads (Smart Life Science) for 2 h at 4°C.. The resin was collected by centrifugation at 500 × *g* for 10 min and loaded onto a gravity flow column. The resin was then washed with 30 column volumes of 20 mM HEPES, 100 mM NaCl, 40 mM imidazole, 100 μM TCEP, 4 μM peptide, 2 mM CaCl_2_, pH 7.4, 0.01% (w/v) LMNG, 0.01% (w/v) GDN and 0.004% (w/v) CHS before bound material was eluted with the same buffer containing 250 mM imidazole. The complexes were concentrated using a 100-kD Amicon Ultra centrifugal filter (Millipore) and loaded onto Superdex 200 10/300 GL column (GE Healthcare) with running buffer containing 20 mM HEPES, 100 mM NaCl, 100 μM TCEP, 4 μM peptide, 2 mM CaCl_2_, pH 7.4, 0.00075% (w/v) LMNG, 0.00025% (w/v) GDN and 0.0002% (w/v) CHS. The monomeric peak fractions were collected and concentrated to 4-6 mg/mL for electron microscopy experiments. Protein concentration was determined by absorbance at 280 nm using a Nanodrop 2000 Spectrophotometer (ThermoFisher).

### Cryo-EM data acquisition

For preparation of cryo-EM grids, 3 μL of the purified MC1R-Gs complex was applied to a glow-discharged holey carbon EM grid (Quantifoil, Au 300 R1.2/1.3) in a Vitrobot chamber (FEI Vitrobot Mark IV). The Vitrobot chamber was set to 100% humidity at 4°C. The sample-coated grids were blotted before plunge-freezing into liquid ethane and stored in liquid nitrogen for data collection. Cryo-EM imaging was performed on a Titan Krios equipped with a Gatan K3 Summit direct electron detector in the Center of Cryo-Electron Microscopy Research Center, Shanghai Institute of Materia Medica, Chinese Academy of Sciences (Shanghai, China). The microscope was operated at 300 kV accelerating voltage, at a nominal magnification of 81,000 ×, corresponding to a pixel size of 1.045 Å. In total, 4,600 movies of α-MSH-MC1R-Gs and 4,600 movies of afamelanotide-MC1R-Gs complexes were obtained at a dose rate of about 22.3 electrons per Å^2^ per second with a defocus range from −0.5 to −3.0 μm. The total exposure time was 3.6 s and the intermediate frames were recorded in 0.1 intervals, resulting in an accumulated dose of 80 electrons per Å^2^ and a total of 36 frames per micrograph. For the SHU9119-MC1R-Gs complex, a total of 6,024 movies were collected with a modified pixel of 1.071 Å. The images were obtained at a dose rate of about 22.3 electrons per Å^2^ per second with a defocus range from −0.5 to −3.0 μm. The total exposure time was 3.2 s and the intermediate frames were recorded in 0.089 intervals, resulting in an accumulated dose of 70 electrons per Å^2^ and a total of 36 frames per micrograph.

### Cryo-EM data processing

Dose-fractioned image stacks were subjected to beam-induced motion correction and dose-weighting using MotionCor2.1^64^. Contrast transfer function parameters for each micrograph were determined by Gctf v1.18^65^. Further data processing was performed with RELION-3.1-beta2^66^.

For the datasets of α-MSH-MC1R-Gs and afamelanotide-MC1R-Gs complexes, particle selection, two-dimensional (2D) classification and three-dimensional (3D) classification were performed on a binned dataset with a pixel size of 2.09 Å. For α-MSH-MC1R-Gs complex, semi-automated selection yielded 4,151,805 particle projections that were subjected to three rounds of reference-free 2D classification to discard false positive particles or particles categorized in poorly defined classes, producing 2,281,404 particle projections for further processing. A well-defined subset of 454,593 particle projections was selected after four rounds of 3D classification and subsequently subjected to 3D refinement, CTF refinement, and Bayesian polishing. The final map has an indicated global resolution of 3.0 Å for α-MSH -MC1R-Gs complex at a Fourier shell correlation of 0.143. For afamelanotide-MC1R-Gs complex, semi-automated selection yielded 3,968,825 particle projections that were subjected to three rounds of reference-free 2D classification to discard false positive particles or particles categorized in poorly defined classes, producing 2,068,327 particle projections for further processing. Two subsets of 814,298 particle projections and 469,220 particle projections were selected after two rounds of 3D classification. Further 3D classifications, focusing on the alignment on the receptor, produced two good subsets of 460,989 particles and 312,962 particles, respectively, which were subsequently subjected to 3D refinement, CTF refinement, and Bayesian polishing. The final maps have an indicated global resolution of 2.7 Å for afamelaonotide-MC1R-Gs-Nb35-scFv16 complex and 2.9 Å for afamelaonotide-MC1R-Gs-scFv16 complex at a Fourier shell correlation of 0.143.

For the datasets of SHU9119-MC1R-Gs complex, particle selection, 2D classification and 3D classification were performed on a binned dataset with a pixel size of 2.142 Å. Semi-automated selection yielded 4,337,394 particle projections that were subjected to three rounds of reference-free 2D classification to discard false positive particles or particles categorized in poorly defined classes, producing 2,162,470 particle projections for further processing. A well-defined subset of 502,722 particle projections was selected after four rounds of 3D classification and subsequently subjected to 3D refinement, CTF refinement, and Bayesian polishing. The final map has an indicated global resolution of 3.1 Å for SHU9119-MC1R-Gs complex at a Fourier shell correlation of 0.143.

The maps were subsequently post-processed in DeepEMhancer^67^. Local resolution was determined using the ResMap with half maps as input maps and surface coloring of the density map was performed using UCSF Chimera^68,69^.

### Model building and refinement

The crystal structure of SHU9119-MC4R complex (PDB: 6W25) was used as the initial model of MC1R for model rebuilding and refinement against the electron microscopy map^26^. The cryo-EM structure of V2R-Gs complex (PDB: 7DW9) was used to generate the initial model of Gs, Nb35 and scFv16^70^. For the structure of afamelanotide-MC1R-Gs and SHU9119-MC1R-Gs complexes, the coordinates of α-MSH-MC1R-Gs complex were used as an initial template. The models were docked into the electron microscopy density maps using UCSF Chimera followed by iterative manual adjustment and rebuilding in Coot^69,71^. Real space refinement and rosetta refinement were performed using ISOLDE and Phenix software package^72,73^. All residues were checked for fitting in electron density, Ramachandran and rotamer restraints. The model statistics was validated using the module ‘comprehensive validation (cryo-EM)’ in Phenix. Finally, MC1R from L36^ECD^ to T308^8.53^, α-MSH (residues Y2-V11), afamelanotide (residues Y2-V11), the full-length SHU9119, and the calcium ion were well defined in the EM maps. However, the fusion partner BRIL, LgBiT, ICL3 of MC1R and AHD of Gαs showed very poor density in the EM maps and were omitted from the final models. Structural figures were prepared in UCSF Chimera, UCSF ChimeraX and PyMOL (https://pymol.org/2/)^74^. The final refinement statistics are provided in **Table S1**.

### Cell culture and transfection

Chinese hamster ovary (CHO-K1) cells were cultured in F12 (Gibco) containing 10% (v/v) fetal bovine serum (FBS, Gibco) at 37°C in 5% CO_2_. Human embryonic kidney 293 cells (HEK293) were maintained in DMEM (Gibco) supplemented with 10% (v/v) FBS, 1 mM sodium pyruvate (Gibco), 100 units/mL penicillin and 100 μg/mL streptomycin at 37°C in 5% CO_2_. For cAMP and MC1R expression level assays, CHO-K1 cells were seeded into 6-well cell culture plates at a density of 5 × 10^5^ cells per well. For whole cell binding assay, HEK293 cells were seeded into 96-well poly-D-lysine-treated cell culture plates at a density of 3 × 10^4^ cells per well. After overnight incubation, cells were transfected with different MC1R constructs using FuGENE^®^ HD transfection reagent (Promega) for cAMP accumulation assay, or Lipofectamine 2000 transfection reagent (Invitrogen) for binding and flow cytometry assays, respectively. Following 24 h culturing, the transfected cells were ready for detection.

### cAMP accumulation assay

α-MSH, afamelanotide and SHU9119 stimulated cAMP accumulation was measured by LANCE Ultra cAMP kit (PerkinElmer). Twenty-four hours post-transfection, CHO-K1 cells were washed and seeded into 384-well microtiter plates at a density of 3,000 cells per well. Then they were incubated with different concentrations of ligands in stimulation buffer (calcium and magnesium free HBSS buffer (Gibco), 5 mM HEPES (Gibco), 0.1% BSA (Abcone) and 0.5 mM IBMX (Abcone)) for 40 min at RT. Eu-cAMP tracer and ULight-anti-cAMP were diluted by cAMP detection buffer and added to the plates separately to terminate the reaction. Plates were incubated at RT for 40 min and read according to the protocol using an EnVision multilabel reader (PerkinElmer) with the emission window ratio of 665 nm over 620 nm. Data were normalized to the wild-type (WT) receptor.

For assessing the effect of calcium ion on cAMP signaling, CHO-K1 cells were dissociated by 0.02% (w/v) EDTA and washed three times with calcium and magnesium free HBSS buffer. Then the cells were resuspended and stimulated with different concentrations of ligands in Ca^2+^ free stimulation buffer consisting of aforementioned stimulation buffer supplemented with 1 mM EGTA, or with additional 1.5 mM CaCl_2_ in Ca^2+^ free stimulation buffer ([Ca^2+^] ~ 0.5 mM)^26^. The rest steps were essentially the same as described above.

### Whole cell binding assay

Radiolabeled ligand binding assays were performed using the whole cell method. In brief, HEK293 cells were harvested 24 h after transfection, washed twice and incubated with blocking buffer (F12 supplemented with 25 mM HEPES and 0.1% BSA, pH 7.4) for 2 h at 37°C. The homogeneous competition binding experiments were conducted by incubating constant concentration of [^125^I]-[Nle^4^, D-Phe^7^]-α-MSH (30 pM, PerkinElmer) with serial dilution of unlabeled ligands [α-MSH (2.38 pM to 5 μM); afamelanotide (2.38 pM to 5 μM); and SHU9119 (2.38 pM to 5 μM)] in binding buffer (DMEM supplemented with 25mM HEPES and 0.1% BSA). For the effect of divalent cations on ligand binding, the cells were incubated with 1 mM EGTA (PBS supplemented with 1% BSA) for 2 h to neutralize divalent cations in medium before addition of 30 pM [^125^I]-[Nle^4^, D-Phe^7^]-α-MSH and varying concentrations of CaCl_2_ and MgCl_2_. The reactions were carried out for 3 h at 37°C and terminated by washing three times with ice-cold PBS. The bound radioactivity was measured with a MicroBeta2 plate counter (PerkinElmer) using a scintillation cocktail (OptiPhase SuperMix, PerkinElmer).

### NanoBiT assay

HEK293A cells (G protein knockout) were seeded into 10-cm plates at a density of 3 × 10^6^ cells per plate and transfected with the plasmid mixture containing 2 μg WT MC1R, 1 μg Gαs-LgBiT, 5 μg WT Gβ or Gβ with mutations F292A and D312A, and 5 μg SmBiT-Gγ using Lipofectamine 3000 transfection reagent (Invitrogen). After 24 h, the cells were transferred to poly-D-lysine coated 96-well plates at a density of 50,000 cells/well and grown overnight before incubation in NanoBiT buffer (calcium and magnesium free HBSS buffer, supplemented with 10 mM HEPES and 0.1% BSA, pH 7.4) in 37℃ for 30 min. Then 10 μL coelentrazine-h (Yeasen Biotech) was added to each well at a working concentration of 5 μM followed by incubation for 2h at room temperature. The luminescence signal was measured using an EnVision plate reader (PerkinElmer) at 30 s interval for 4 min as baseline, and then read for 10 min after addition of ligand. Data were corrected to baseline measurements and then the vehicle control to determine ligand-induced changes in response. Dose-response values were obtained from the area-under-the-curve of elicited responses by each ligand.

### Receptor expression

Membrane expression of MC1R was determined by flow cytometry to detect the N-terminal Flag tag on the WT and mutated receptor constructs transiently expressed in CHO-K1 cells. Briefly, approximately 2 × 10^5^ cells were blocked with PBS containing 5% BSA (w/v) at RT for 15 min, and then incubated with 1:300 anti-Flag primary antibody (diluted with PBS containing 5% BSA, Sigma) at RT for 1 h. The cells were then washed three times with PBS containing 1% BSA (w/v) followed by 1 h incubation with 1:1000 anti-mouse Alexa Fluor 488 conjugated secondary antibody (diluted with PBS containing 5% BSA, Invitrogen) at RT in the dark. After washing three times, cells were resuspended in 200 μL PBS containing 1% BSA for detection by NovoCyte (Agilent) utilizing laser excitation and emission wavelengths of 488 nm and 530 nm, respectively. For each sample, 20,000 cellular events were collected, and the total fluorescence intensity of positive expression cell population was calculated. Data were normalized to WT receptor and parental CHO-K1 cells.

### Data analysis

Dose-response data were analyzed using Prism 8 (GraphPad). Non-linear curve fit was performed using a three-parameter logistic equation [log (agonist *vs*. response)]. All data are presented as means ± S.E.M. of at least three independent experiments. Statistical significance was determined by Dunnett’s test.

## ACKNOWLEDGEMENTS

The Cryo-EM data were collected at Cryo-Electron Microscopy Research Center, Shanghai Institute of Material Medica, Chinese Academy of Sciences. We thank the staff of the National Center for Protein Science (Shanghai) Electron Microscopy facility for instrument support. This work was partially supported by the Ministry of Science and Technology (China) grants 2018YFA0507002 (H.E.X.) and 2018YFA0507000 (M.-W.W.); National Natural Science Foundation of China 31770796 (Y.J.), 81872915 (M.-W.W.), 82073904 (M.-W.W.), 81773792 (D.Y.), and 81973373 (D.Y.); National Science and Technology Major Project of China – Key New Drug Creation and Manufacturing Program 2018ZX09711002-002-002 (Y.J.), 2018ZX09735–001 (M.-W.W.), and 2018ZX09711002–002–005 (D.Y.); the Shanghai Municipal Science and Technology Commission Major Project 2019SHZDZX02 (H.E.X.); the CAS Strategic Priority Research Program XDB37030103 (H.E.X.); and Novo Nordisk-CAS Research Fund grant NNCAS-2017–1-CC (D.Y.).

## AUTHOR CONTRIBUTIONS

S.M. designed the expression constructs, optimized and purified the MC1R-Gs protein complexes, prepared the cryo-EM grids, collected the cryo-EM images, performed the structure determination and model building, participated in the preparation of the constructs for functional assays, analyzed the structures, prepared the figures and wrote the manuscript; Y.C. prepared the constructs for functional assays, performed the cAMP accumulation and MC1R surface expression assays, participated in the figure preparation; A.D. performed the whole cell binding assay; W.Y. designed the Gαs construct and participated in the cryo-EM grids preparation. J.G. and F.Z. participated the data analysis and manuscript editing; D.Y. performed the data analysis and participated in the manuscript editing; Y.J. participated in project supervision and manuscript editing; M.-W.W. oversaw the work of C.Y., A.D. and D.Y. and participated in the manuscript writing; H.E.X. conceived and supervised the project, analyzed the structures, and wrote the manuscript with input from all co-authors.

