## Supplementary Fig. S1-S7 and Table S1-S6 for "Structural mechanism of calcium-mediated hormone recognition and Gβ interaction by the human melanocortin-1 receptor"

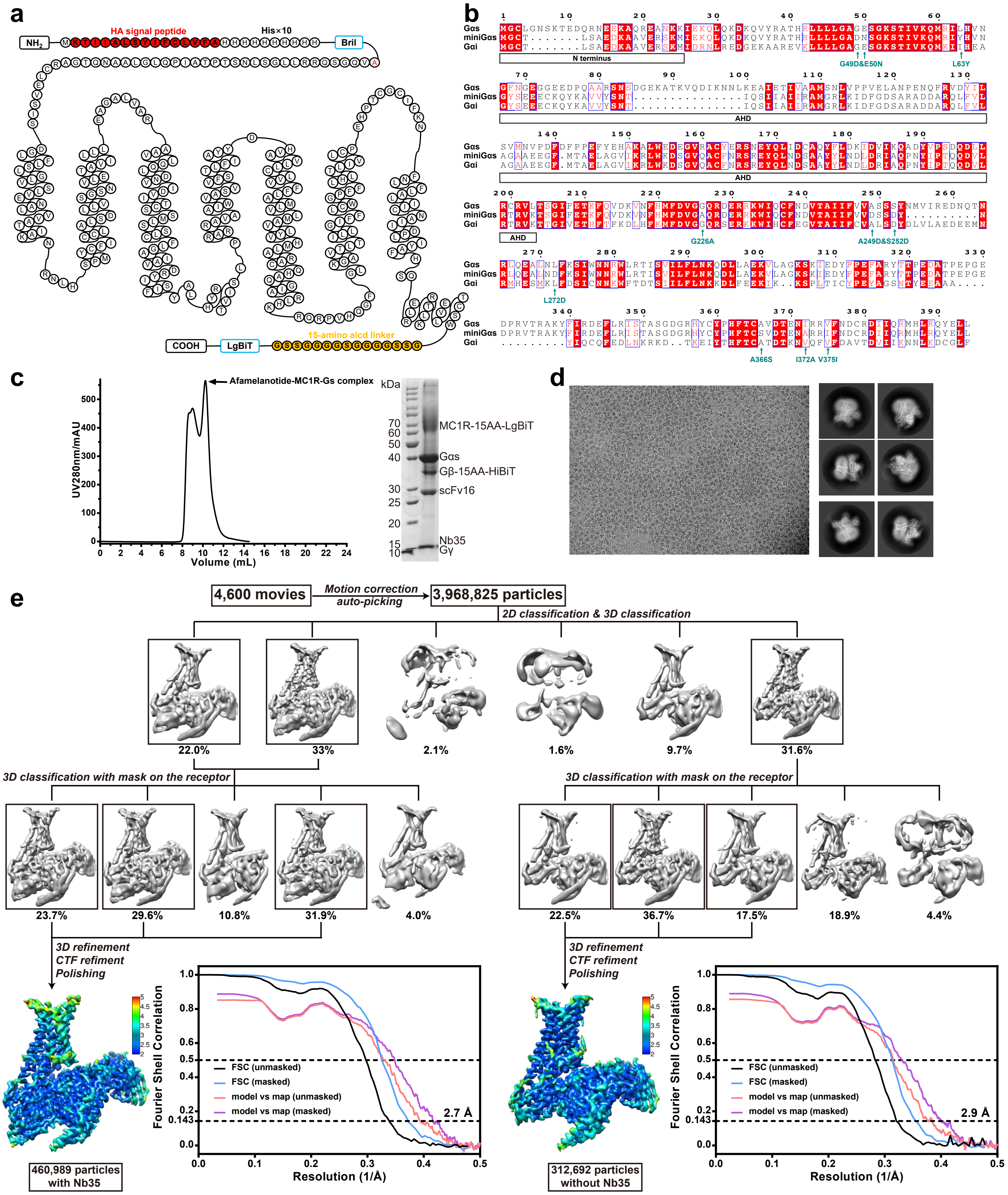


**Fig. S1 Constructs of MC1R and Gαs and cryo-EM structure determination of the afamelanotide-MC1R-Gs complex. a** Schematic diagram of the human MC1R construct used in this study. **b** Sequence alignment of miniG αs used in this study with human Gαs and human Gαi. The replacement of the N terminus and AHD of Gα are marked by black frames and the mutations are indicated by teal arrows. **c** The size-exclusion chromatography elution profile (left panel) and SDS-PAGE analysis (right panel) of the afamelanotide-MC1R-Gs complex. **d** Representative cryo-EM micrograph and 2D class averages showing distinct secondary structure features from different views (scale bar: 5 nm) of the afamelanotide-MC1R-Gs complex. **e** The flowchart of cryo-EM data analysis, the cryo-EM maps colored by local resolution and the ‘Gold-standard’ Fourier shell correlation curves of the afamelanotide-MC1R-Gs complex.

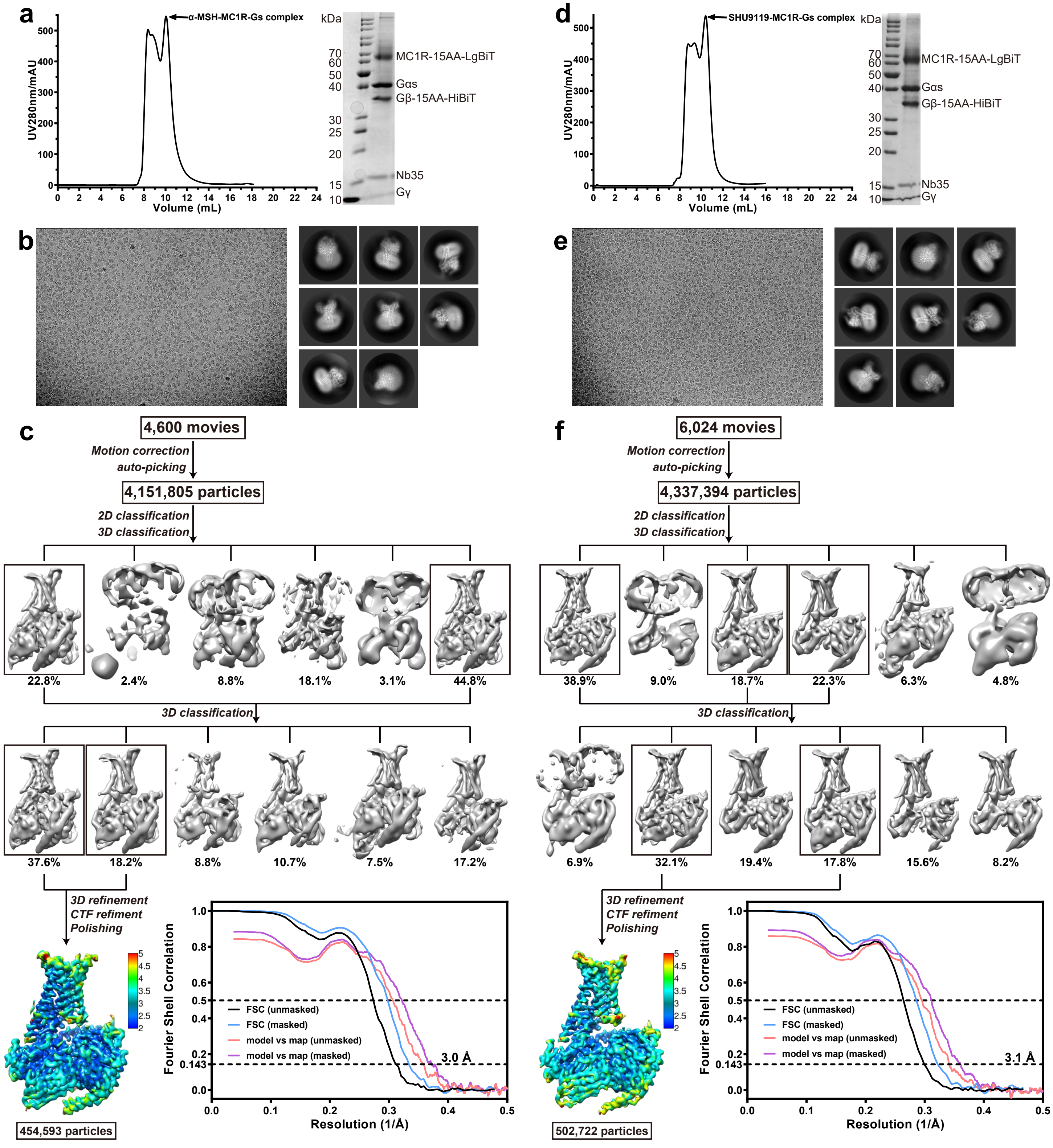


**Fig. S2 Cryo-EM structure determination of the α-MSH-MC1R-Gs and SHU9119-MC1R-Gs complexes.** **a**, **d** Size-exclusion chromatography elution profiles (left panel) and SDS-PAGE analysis (right panel) of the α-MSH-MC1R-Gs complex (**a**) and SHU9119-MC1R-Gs complex (**d**). **b**, **e** Representative cryo-EM micrographs and 2D class averages showing distinct secondary structure features from different views (scale bar: 5 nm) of the α-MSH-MC1R-Gs complex (**b**) and SHU9119-MC1R-Gs complex (**e**). **c**, **f** Flowchart of cryo-EM data analysis, cryo-EM maps colored by local resolution and 'Gold-standard' Fourier shell correlation curves of the α-MSH-MC1R-Gs complex (**c**) and SHU9119-MC1R-Gs complex (**f**).

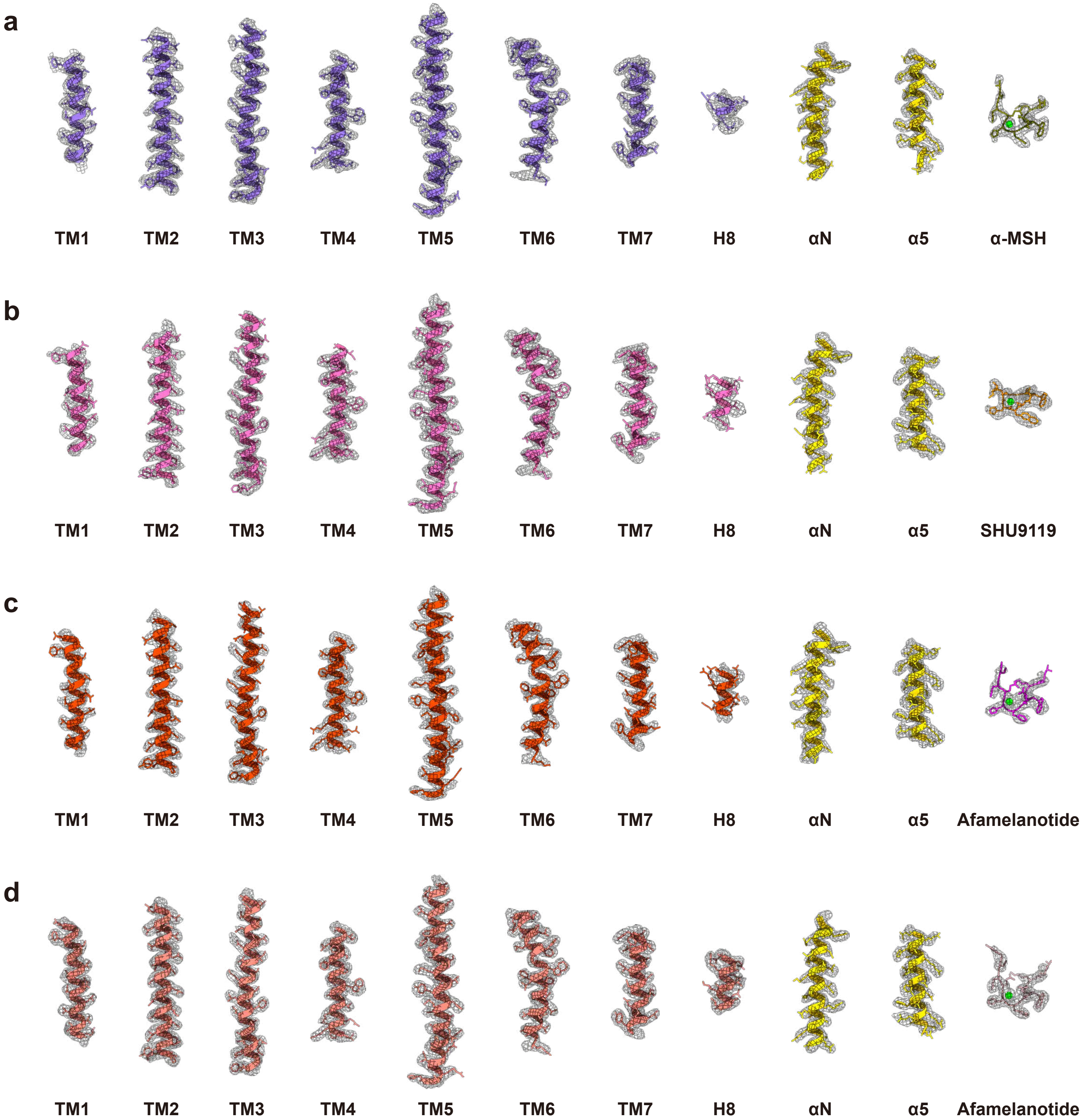


**Fig. S3 Atomic resolution models of the MC1R-Gs complexes.** EM density maps and models of the α-MSH-MC1R-Gs complex (**a**), SHU9119-MC1R-Gs complex (**b**), afamelanotide-MC1R-Gs-Nb35-scFv16 complex (**c**), and afamelanotide-MC1R-Gs-scFv16 complex (**d**). All transmembrane helices of MC1R, α5 helix and αN helix of Gαs subunit, α-MSH, SHU9119, afamelanotide and calcium ion are shown. The EM densities are shown at 0.05 threshold.

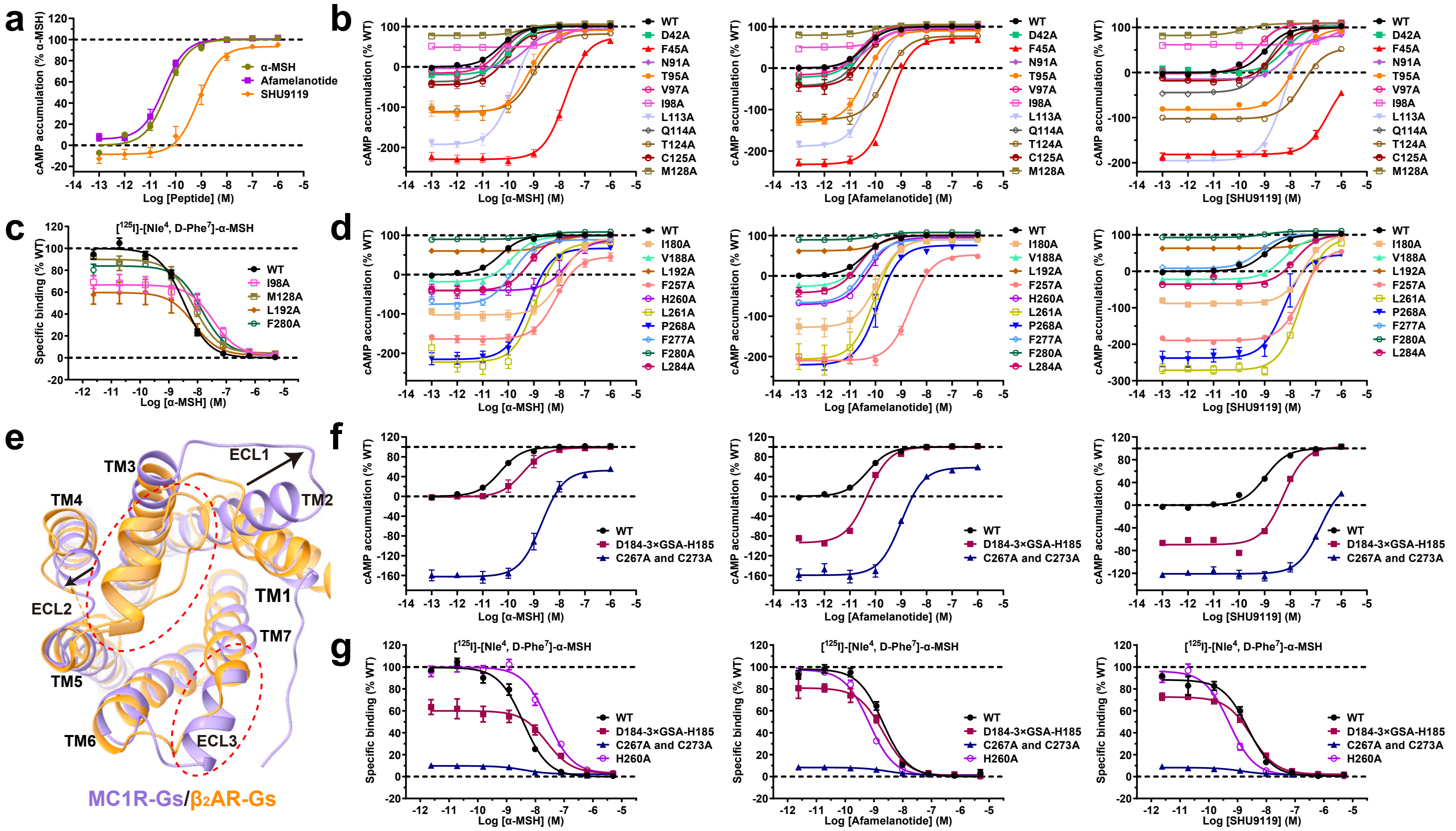

**Fig. S4 cAMP accumulation and melanocortin peptide ligand binding of different MC1R constructs.** **a** Dose-response curves of cAMP accumulation of the WT MC1R stimulated by α-MSH, afamelanotide and SHU9119. Data are presented as means ± S.E.M. of three independent experiments performed in quadruplicate. **b**, **d** Dose-response curves of cAMP accumulation of WT MC1R and MC1R with mutations in the orthosteric binding pocket stimulated by α-MSH, afamelanotide and SHU9119. Data are presented as means ± S.E.M. of three independent experiments performed in quadruplicate. **c** Effects of mutations in the orthosteric binding pocket on α-MSH binding ability assessed by a radiolabeled ligand binding assay. Competition curves of representative mutants with marked changes in the basal activity compared to WT MC1R are shown. **e** Structural comparison of the extracellular side of MC1R with $β_2$AR (PDB: 3SN6). The differences of extracellular loops are marked by black arrows. TM2 and ECL1 of MC1R move outwards to form a large opening of TMD. The long ECL2 of $β_2$AR forms a lid covering the extracellular side of TMD, which is not observed in MC1R. **f** Dose-response curves of cAMP accumulation of WT MC1R and MC1R with mutations in ECL2 and ECL3 stimulated by α-MSH, afamelanotide and SHU9119. Data are presented as means ± S.E.M. of three independent experiments performed in quadruplicate. **g** Competition curves of WT MC1R, MC1R with H260A mutation and MC1R with mutations in ECL2 and ECL3 are shown. The affinity of three peptides were assessed by a radiolabeled ligand binding assay.

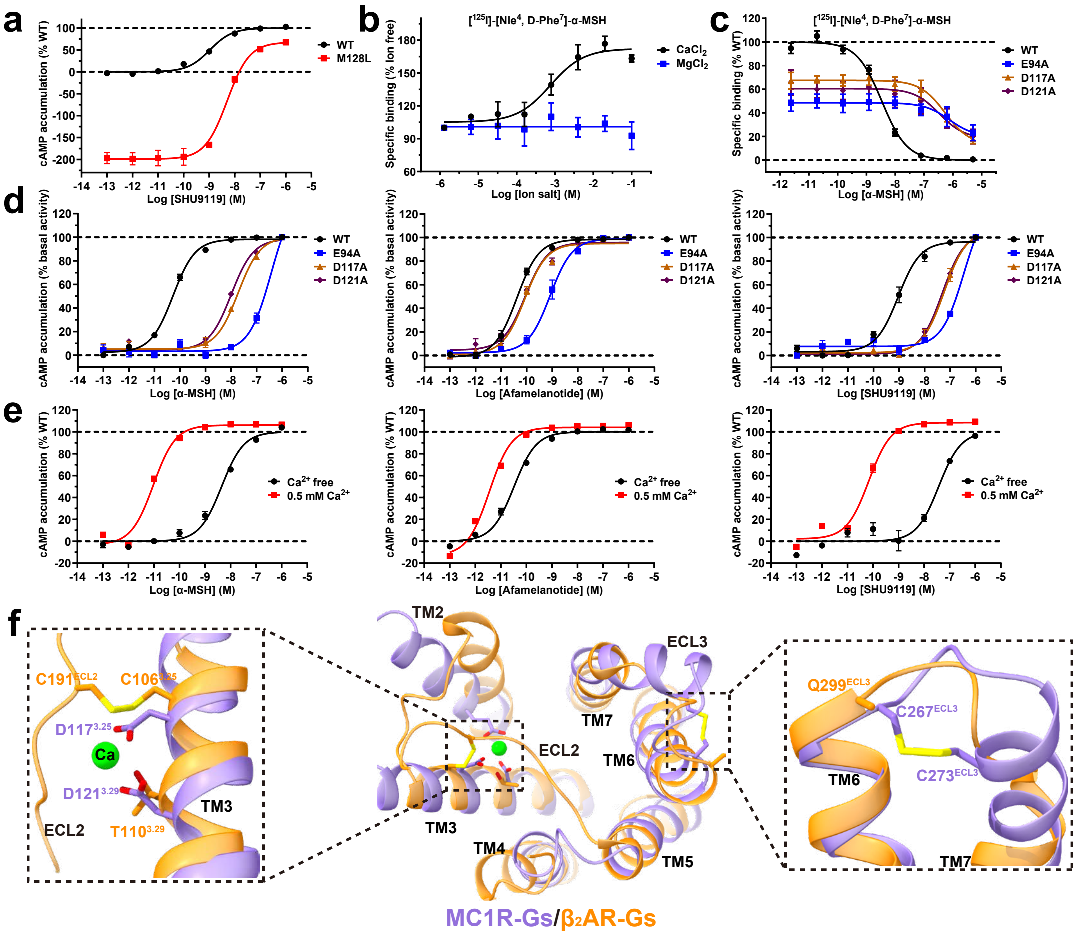

**Fig. S5 Differential activities of melanocortin peptide ligands on melanocortin receptors and the role of calcium ion on ligand binding. a** Dose-response curves of cAMP accumulation of the WT and mutant (M128L) MC1Rs stimulated by SHU9119. Data are presented as means ± S.E.M. of three independent experiments performed in quadruplicate. **b** Effects of different metal ions on α-MSH-binding ability compared to WT MC1R assessed by a radiolabeled ligand binding assay. **c** Effects of mutations of MC1R on peptide-binding ability assessed by a radiolabeled ligand binding assay. Competition curves of MC1R mutants compared to WT MC1R are shown. **d** Dose-response curves of cAMP accumulation of WT MC1R and MC1R with mutations in the calcium-binding pocket stimulated by α-MSH, afamelanotide and SHU9119. Data are presented as means ± S.E.M. of three independent experiments performed in quadruplicate. **e** Dose-response curves of cAMP accumulation of WT MC1R stimulated by α-MSH, afamelanotide and SHU9119 under the condition of $Ca^{2+}$ free or 0.5 mM $Ca^{2+}$. Data are presented as means ± S.E.M. of three independent experiments performed in quadruplicate. **f** Structural comparison of MC1R with $\beta_2$AR (PDB: 3SN6) from the top view (middle panel). The canonical disulfide bond between TM3 and ECL2 of $\beta_2$AR (left panel) and the unique disulfide bond between ECL3 of MC1R (right panel) are emphasized. The alignment was based on the structures of MC1R and $\beta_2$AR.

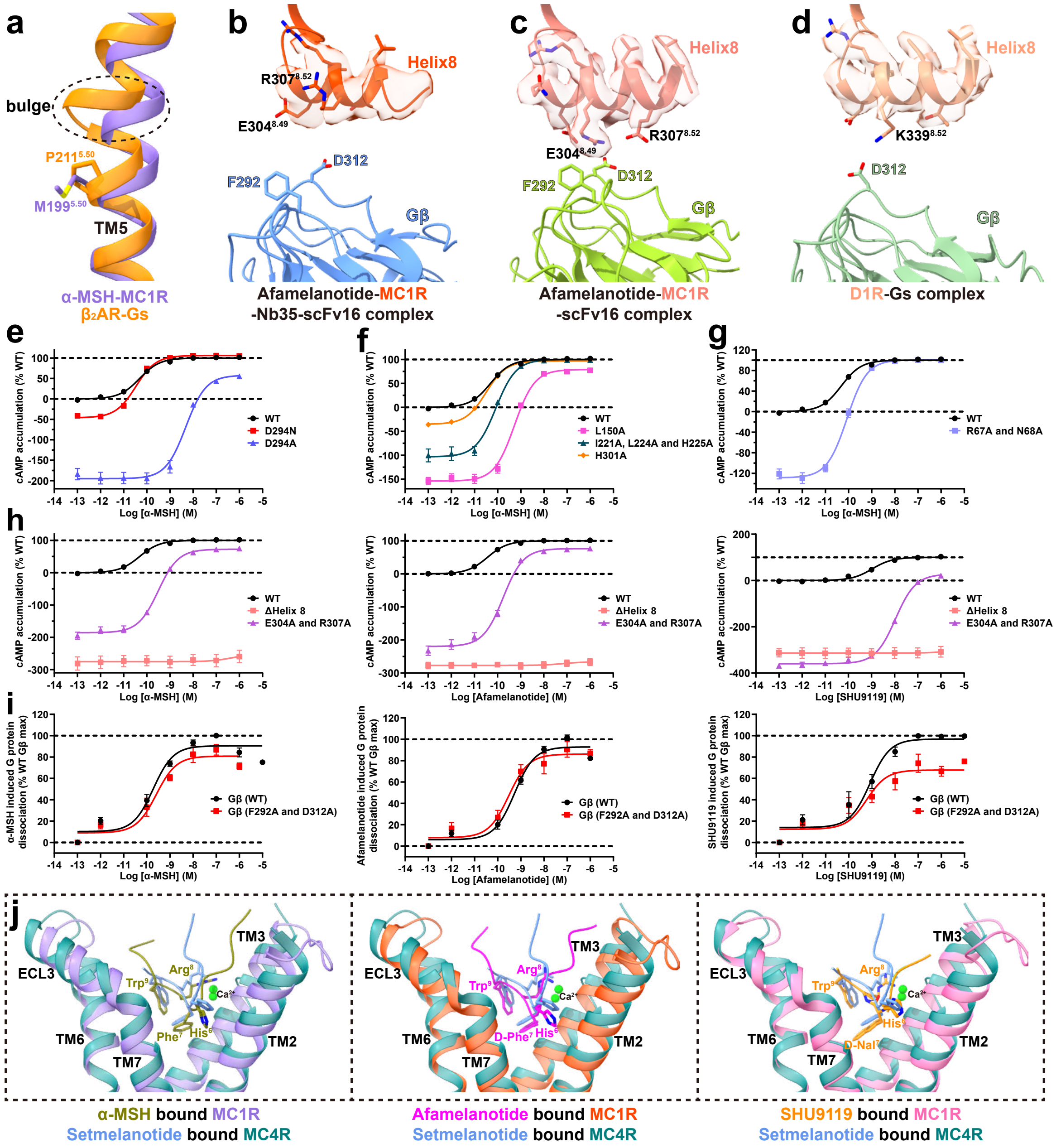


**Fig. S6 The mechanism of MC1R activation and Gs coupling. a** Structural comparison between $M^{5.50}$ of MC1R (medium purple) and $P^{5.50}$ of β2AR (PDB: 3SN6, dark orange). The bulge of TM5 is not observed in MC1R. The alignment was based on the structures of MC1R and β2AR. **b-d** The interactions between helix 8 and Gβ in the afamelanotide-MC1R-Gs-Nb35-scFv16 complex (**b**), afamelanotide-MC1R-Gs-scFv16 complex (**c**) and D1R-Gs complex (**d**, PDB: 7JVP). The EM densities of helix 8 of MC1R and D1R are shown. **e-g** Dose-response curves of cAMP accumulation of the WT and mutated MC1Rs stimulated by α-MSH. Data are presented as means ± S.E.M. of three independent experiments performed in quadruplicate. **h** Dose-response curves of cAMP accumulation of WT MC1R and MC1R with mutations in helix 8 stimulated by α-MSH, afamelanotide and SHU9119. Data are presented as means ± S.E.M. of three independent experiments performed in quadruplicate. **i** α-MSH, afamelanotide and SHU9119 induced G protein activation of WT Gβ and Gβ with mutations in the corresponding residues using NanoBiT assay. Data are presented as means ± S.E.M. of three independent experiments performed in quadruplicate. **j** Structural comparison of the orthosteric binding pockets among α-MSH bound MC1R complex, afamelanotide bound MC1R complex, SHU9119 bound MC1R complex and setmelanotide bound MC4R complex (PDB: 7AUE). The alignment was based on the structures of MC1R and MC4R.

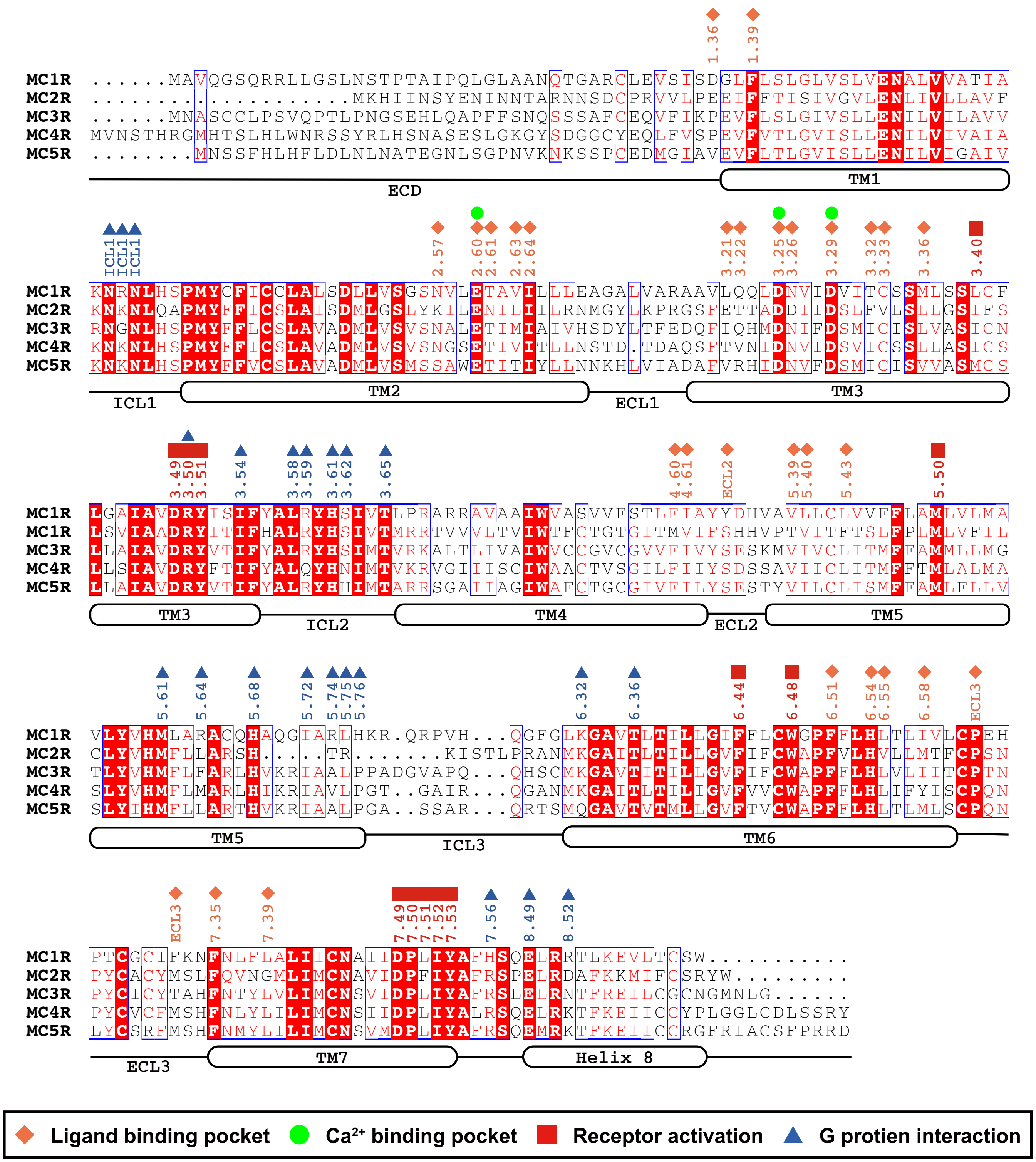


**Fig. S7 Sequence alignment of melanocortin receptors.** Superscripts refer to the Ballesteros-Weinstein numbers.

**Table S1 Cryo-EM data collection, model refinement and validation statistics.**

| MC1R-Gs complex | α-MSH-MC1R-Gs-Nb35 | Afamelanotide-MC1R-Gs-scFv16 | Afamelanotide-MC1R-Gs-Nb35-scFv16 | SHU9119-MC1R-Gs-Nb35 |
|---|---|---|---|---|
| **Data collection and processing** | | | | |
| Magnification | 81,000 | 81,000 | 81,000 | 81,000 |
| Voltage (kV) | 300 | 300 | 300 | 300 |
| Electron exposure ($e^{-}$/Å$^2$) | 80 | 80 | 80 | 70 |
| Defocus range (μm) | -0.5 to -3.0 | -0.5 to -3.0 | -0.5 to -3.0 | -0.5 to -3.0 |
| Pixel size (Å) | 1.045 | 1.045 | 1.045 | 1.071 |
| Symmetry imposed | C1 | C1 | C1 | C1 |
| Initial particle projections (no.) | 4,151805 | 3,968,825 | 3,968,825 | 4,337,394 |
| Final particle projections (no.) | 454,593 | 312,962 | 460,989 | 502,722 |
| Map resolution (Å) | 3.0 | 2.9 | 2.7 | 3.1 |
| FSC threshold | 0.143 | 0.143 | 0.143 | 0.143 |
| Map resolution range (Å) | 2.0-5.0 | 2.0-5.0 | 2.0-5.0 | 2.0-5.0 |
| **Refinement** | | | | |
| Initial model used (PDB code) | PDB: 6W25 | PDB: 6W25 | PDB: 6W25 | PDB: 6W25 |
| Model resolution (Å) | 3.1 | 3.0 | 2.9 | 3.2 |
| FSC threshold | 0.5 | 0.5 | 0.5 | 0.5 |
| Model resolution range (Å) | 50-3.0 | 50-2.9 | 50-2.7 | 50-3.1 |
| Map sharpening *B* factor (Å$^2$) | -116.20 | -80.97 | -76.84 | -122.70 |
| Model composition | | | | |
| Non-hydrogen atoms | 8048 | 8882 | 9850 | 8032 |
| Protein residues | 1030 | 1138 | 1265 | 1030 |
| Lipids | 0 | 0 | 0 | 0 |
| *B* factors (Å$^2$) | | | | |
| Protein | 111.94 | 48.91 | 89.69 | 129.70 |
| RMSD | | | | |
| Bond lengths (Å) | 0.009 | 0.010 | 0.010 | 0.009 |
| Bond angles (°) | 1.066 | 1.100 | 1.043 | 1.093 |
| Validation | | | | |
| MolProbity score | 1.40 | 1.50 | 1.46 | 1.22 |
| Clashscore | 3.86 | 4.64 | 3.88 | 1.69 |
| Rotamer outliers (%) | 0.00 | 0.00 | 0.00 | 0.00 |
| Ramachandran plot | | | | |
| Favored (%) | 96.55 | 96.14 | 95.89 | 95.74 |
| Allowed (%) | 3.45 | 3.86 | 4.11 | 4.26 |
| Disallowed (%) | 0.00 | 0.00 | 0.00 | 0.00 |
| **PDB accession number** | 7F4D | 7F4F | 7F4H | 7F4I |
| **EMDB accession number** | EMD-31448 | EMD-31449 | EMD-31452 | EMD-31453 |

**Table S2 Interactions of α-MSH with MC1R.**

| α-MSH | MC1R | Interactions |
|---|---|---|
| $Tyr^{M2}$ | $A110^{3.18}$ | Hydrophobic interactions |
| | $Q114^{3.22}$ | |
| $Ser^{M3}$ | $Q114^{3.22}$ | Hydrogen bonds |
| $Met^{M4}$ | $E94^{2.60}$ | Hydrophobic interactions |
| | $V97^{2.63}$ | |
| | $I98^{2.64}$ | |
| | $L113^{3.21}$ | |
| | $D117^{3.25}$ | |
| | $I120^{3.28}$ | |
| $Glu^{M5}$ | $E94^{2.60}$ | Van der Waals force |
| | $D117^{3.25}$ | |
| $His^{M6}$ | $F45^{1.39}$ | π–π interaction |
| | $N91^{2.57}$ | Hydrophobic interactions |
| | $E94^{2.60}$ | |
| | $T95^{2.61}$ | |
| | $L284^{7.39}$ | |
| $Phe^{M7}$ | $E94^{2.60}$ | Hydrophobic interactions |
| | $D121^{3.29}$ | |
| | $T124^{3.32}$ | |
| | $M128^{3.36}$ | |
| | $F257^{6.51}$ | |
| | $F280^{7.35}$ | |
| $Arg^{M8}$ | $D117^{3.25}$ | Hydrogen bonds |
| | $N118^{3.26}$ | |
| | $D121^{3.29}$ | Hydrophobic interactions |
| | $I180^{4.61}$ | |
| | $Y183^{4.64}$ | |
| | $F280^{7.35}$ | |
| $Trp^{M9}$ | $Y183^{ECL2}$ | Hydrogen bonds |
| | $H260^{6.54}$ | |
| | $F179^{4.60}$ | Hydrophobic interactions |
| | $V188^{5.39}$ | |
| | $L189^{5.40}$ | |
| | $L192^{5.43}$ | |
| | $L261^{6.55}$ | |
| | $I264^{6.58}$ | |
| $Gly^{M10}$ | $F277^{ECL3}$ | Hydrophobic interactions |
| | $F280^{7.35}$ | |
| $Pro^{M12}$ | $I264^{6.58}$ | Hydrophobic interactions |
| | $P268^{ECL3}$ | |
| | $F277^{ECL3}$ | |
| $Val^{M13}$ | $D184^{ECL2}$ | Hydrophobic interactions |

Superscripts refer to the Ballesteros-Weinstein numbers and residues within 4 Å are shown.

**Table S3 Interactions of afamelanotide with MC1R.**

| Afamelanotide | MC1R | Interactions |
|---|---|---|
| $Tyr^{F2}$ | $L113^{3.21}$ | Hydrophobic interactions |
| $Nle^{F4}$ | $V97^{2.63}$<br>$I98^{2.64}$<br>$D117^{3.25}$ | Hydrophobic interactions |
| $Glu^{F5}$ | $E94^{2.60}$<br>$D117^{3.25}$ | Van der Waals force |
| $His^{F6}$ | $D42^{1.36}$ | Hydrogen bond |
| | $F45^{1.39}$ | π–π interaction |
| | $E94^{2.60}$<br>$T95^{2.61}$<br>$F280^{7.35}$<br>$L284^{7.39}$ | Hydrophobic interactions |
| $D\text{-}Phe^{F7}$ | $E94^{2.60}$ | Hydrogen bonds |
| | $D121^{3.29}$<br>$T124^{3.32}$<br>$C125^{3.33}$<br>$M128^{3.36}$<br>$F257^{6.51}$<br>$L284^{7.39}$ | Hydrophobic interactions |
| $Arg^{F8}$ | $D117^{3.25}$<br>$D121^{3.29}$ | Hydrogen bonds |
| | $I180^{4.61}$<br>$Y183^{ECL2}$<br>$F280^{7.35}$ | Hydrophobic interactions |
| $Trp^{F9}$ | $Y183^{ECL2}$ | Hydrogen bond |
| | $I180^{4.61}$<br>$L189^{5.40}$<br>$H260^{6.54}$<br>$L261^{6.55}$<br>$I264^{6.58}$ | Hydrophobic interactions |
| $Gly^{F10}$ | $H260^{6.55}$ | Hydrogen bond |
| | $F277^{ECL3}$<br>$F280^{7.35}$ | Hydrophobic interactions |
| $Pro^{F12}$ | $P268^{ECL3}$<br>$F277^{ECL3}$ | Hydrophobic interactions |
| $Val^{F13}$ | $P268^{ECL3}$ | Hydrophobic interactions |

Superscripts refer to the Ballesteros-Weinstein numbers and residues within 4 Å are shown.

**Table S4 Interactions of SHU9119 with MC1R.**

| SHU9119 | MC1R | Interactions |
|---|---|---|
| $Nle^{U1}$ | $V97^{2.63}$ | Hydrophobic interactions |
| | $I98^{2.64}$ | |
| | $D117^{3.25}$ | |
| $Asp^{U2}$ | $E94^{2.60}$ | Van der Waals force |
| | $I98^{2.64}$ | |
| $His^{U3}$ | $F45^{1.39}$ | π–π interaction |
| | $T95^{2.61}$ | Hydrogen bond |
| | $E94^{2.60}$ | Hydrophobic interactions |
| | $I98^{2.64}$ | |
| | $F280^{7.35}$ | |
| | $L284^{7.39}$ | |
| $D\text{-}Nal^{U4}$ | $E94^{2.60}$ | Hydrogen bond |
| | $D121^{3.29}$ | Hydrophobic interactions |
| | $C125^{3.33}$ | |
| | $M128^{3.36}$ | |
| | $F257^{6.51}$ | |
| | $L284^{7.39}$ | |
| $Arg^{U5}$ | $D121^{3.29}$ | Hydrogen bond |
| | $I180^{4.61}$ | Hydrophobic interactions |
| | $Y183^{ECL2}$ | |
| | $F280^{7.35}$ | |
| $Trp^{U6}$ | $Y183^{ECL2}$ | Hydrogen bond |
| | $V188^{5.39}$ | Hydrophobic interactions |
| | $L189^{5.40}$ | |
| | $L192^{5.43}$ | |
| | $H26^{6.54}$ | |
| | $L261^{6.55}$ | |
| | $I264^{6.58}$ | |
| $Lys^{U7}$ | $F280^{7.35}$ | Hydrophobic interactions |

Superscripts refer to the Ballesteros-Weinstein numbers and residues within 4 Å

**Table S5 cAMP accumulation and membrane expression of WT and mutant MC1Rs.**

| MC1R | Expression (% WT) | α-MSH | | | Afamelanotide | | | SHU9119 | | |
|---|---|---|---|---|---|---|---|---|---|---|
| | | $pEC_{50}$ | $\Delta pEC_{50}$ | $E_{max}$ (% WT) | $pEC_{50}$ | $\Delta pEC_{50}$ | $E_{max}$ (% WT) | $pEC_{50}$ | $\Delta pEC_{50}$ | $E_{max}$ (% WT) |
| **WT** | 100 | 10.31 ± 0.03 | 0 | 100.01 ± 0.75 | 10.44 ± 0.03 | 0 | 99.94 ± 0.66 | 9.00 ± 0.06 | 0 | 99.41 ± 1.79 |
| **D42A** | 113.55 ± 7.43 | 9.90 ± 0.08 | -0.49 ± 0.07** | 100.70 ± 2.66 | 10.60 ± 0.10 | -0.07 ± 0.06 | 100.52 ± 3.03 | 8.17 ± 0.06**** | -1.35 ± 0.04**** | 103.87 ± 2.59 |
| **F45A** | 103.95 ± 5.22 | 7.77 ± 0.07**** | -2.42 ± 0.06**** | 73.68 ± 8.6*** | 9.46 ± 0.05**** | -0.87 ± 0.04**** | 72.17 ± 4.29**** | 6.52 ± 0.13**** | -2.36 ± 0.09**** | -3.19 ± 15.84**** |
| **R67A and N68A** | 112.84 ± 3.86 | 10.07 ± 0.05 | -0.34 ± 0.10 | 100.66 ± 3.31 | N.D. | N.D. | N.D. | N.D. | N.D. | N.D. |
| **N91A** | 135.10 ± 5.23** | 9.27 ± 0.08**** | -0.98 ± 0.05**** | 93.51 ± 2.33 | 10.24 ± 0.05 | -0.22 ± 0.03** | 96.01 ± 1.24 | 8.14 ± 0.08**** | -1.23 ± 0.03**** | 80.91 ± 3.13 |
| **E94A** | 134.13 ± 4.27** | 6.21 ± 0.43**** | -3.81 ± 0.04**** | 104.58 ± 10.99 | 8.91 ± 0.05**** | -1.55 ± 0.01**** | 95.52 ± 1.40 | 6.50 ± 0.23**** | -2.37 ± 0.11**** | 103.15 ± 4.95 |
| **T95A** | 99.34 ± 6.25 | 9.10 ± 0.11**** | -1.28 ± 0.10**** | 93.48 ± 4.44 | 10.38 ± 0.06 | -0.21 ± 0.02* | 90.21 ± 3.17 | 7.85 ± 0.05**** | -1.67 ± 0.04**** | 97.09 ± 3.74 |
| **V97A** | 104.66 ± 6.53 | 10.27 ± 0.08 | -0.13 ± 0.06 | 103.23 ± 2.56 | 10.62 ± 0.07 | -0.06 ± 0.06 | 104.32 ± 1.95 | 9.31 ± 0.03 | -0.20 ± 0.04 | 109.17 ± 0.92 |
| **I98A** | 114.26 ± 7.03 | 7.86 ± 0.07**** | -2.25 ± 0.12**** | 102.25 ± 1.68 | 9.60 ± 0.14**** | -0.71 ± 0.04**** | 98.25 ± 1.89 | 6.68 ± 0.21**** | -2.34 ± 0.11**** | 98.95 ± 5.52 |
| **L113A** | 88.60 ± 4.44 | 9.77 ± 0.05* | -0.28 ± 0.12 | 101.93 ± 4.48 | 10.13 ± 0.03 | -0.20 ± 0.01* | 104.33 ± 2.47 | 8.21 ± 0.03*** | -0.44 ± 0.01* | 109.21 ± 3.87 |
| **Q114A** | 97.51 ± 8.19 | 10.25 ± 0.06 | 0.04 ± 0.11 | 101.55 ± 2.11 | 10.60 ± 0.04 | 0.12 ± 0.13 | 102.54 ± 1.25 | 8.84 ± 0.03 | 0.05 ± 0.13 | 104.43 ± 1.69 |
| **D117A** | 124.47 ± 7.41 | 7.61 ± 0.08**** | -2.40 ± 0.06**** | 100.75 ± 2.13 | 10.02 ± 0.22** | -0.24 ± 0.05** | 108.45 ± 1.89 | 7.13 ± 0.27**** | -1.82 ± 0.1**** | 109.74 ± 7.15 |
| **D121A** | 143.35 ± 10**** | 8.00 ± 0.07**** | -2.14 ± 0.05**** | 101.52 ± 2.15 | 10.12 ± 0.08 | -0.17 ± 0.03 | 102.04 ± 1.51 | 7.26 ± 0.07**** | -1.53 ± 0.15**** | 103.88 ± 2.02 |
| **T124A** | 94.83 ± 6.27 | 9.00 ± 0.10**** | -1.49 ± 0.07**** | 81.58 ± 6.13 | 9.61 ± 0.08**** | -1.07 ± 0.08**** | 77.16 ± 4.29**** | 7.52 ± 0.04**** | -2.00 ± 0.04**** | 54.35 ± 2.60**** |
| **C125A** | 95.84 ± 5.51 | 10.05 ± 0.10 | -0.35 ± 0.07 | 91.61 ± 3.84 | 10.43 ± 0.14 | -0.24 ± 0.06*** | 94.22 ± 4.68 | 8.61 ± 0.03 | -0.91 ± 0.04**** | 98.51 ± 1.38 |
| **M128A** | 154.18 ± 10.72**** | 9.41 ± 0.15**** | -0.63 ± 0.30**** | 105.93 ± 1.21 | 10.09 ± 0.12* | -0.24 ± 0.03** | 105.32 ± 0.86 | 9.79 ± 0.16**** | 1.10 ± 0.19**** | 108.23 ± 1.19 |
| **M128L** | 113.20 ± 9.40 | N.D. | N.D. | N.D. | N.D. | N.D. | N.D. | 8.56 ± 0.17 | -0.5 ± 0.02** | 57.41 ± 14.88**** |
| **L150A** | 105.88 ± 4.45 | 9.27 ± 0.05**** | -1.14 ± 0.07**** | 79.11 ± 3.90** | N.D. | N.D. | N.D. | N.D. | N.D. | N.D. |
| **I180A** | 112.34 ± 1.39 | 7.91 ± 0.10**** | -2.73 ± 0.12**** | 93.51 ± 8.57 | 9.94 ± 0.13*** | -0.71 ± 0.05**** | 90.40 ± 7.96 | 7.46 ± 0.03**** | -2.06 ± 0.04**** | 100.87 ± 2.97 |
| **D184A-3×GSA-H185A** | 121.35 ± 5.72 | 9.42 ± 0.12**** | -1.68 ± 0.16**** | 98.46 ± 3.60 | 10.31 ± 0.04 | 0.01 ± 0.01 | 100.09 ± 2.11 | 8.27 ± 0.06*** | -0.55 ± 0.01*** | 102.44 ± 3.98 |
| **V188A** | 90.08 ± 2.83 | 9.87 ± 0.09 | -0.26 ± 0.08 | 97.77 ± 3.02 | 10.54 ± 0.05 | 0.02 ± 0.15 | 99.92 ± 1.43 | 8.25 ± 0.13*** | -0.47 ± 0.11** | 94.22 ± 5.77 |
| **L192A** | 104.22 ± 9.34 | 8.29 ± 0.15**** | -1.84 ± 0.10**** | 98.96 ± 2.16 | 9.80 ± 0.13**** | -0.66 ± 0.05**** | 100.73 ± 1.45 | 7.17 ± 0.21**** | -1.60 ± 0.02**** | 102.31 ± 4.44 |
| **I221A, L224A and H225A** | 117.48 ± 11.43 | 10.08 ± 0.07 | -0.33 ± 0.08 | 99.88 ± 3.91 | N.D. | N.D. | N.D. | N.D. | N.D. | N.D. |
| **F257A** | 74.71 ± 5.23 | 8.13 ± 0.08**** | -2.18 ± 0.10**** | 45.81 ± 6.79**** | 8.63 ± 0.07**** | -1.86 ± 0.09**** | 52.19 ± 6.20**** | 7.53 ± 0.06**** | -1.49 ± 0.05**** | 62.82 ± 7.17**** |
| **H260A** | 93.76 ± 6.34 | 7.71 ± 0.27**** | -2.40± 0.22**** | 89.12 ± 14.85 | 10.29 ± 0.09 | -0.14 ± 0.05 | 95.04 ± 3.98 | N.D. | N.D. | N.D. |
| **L261A** | 86.01 ± 4.46 | 8.94 ± 0.10**** | -1.05 ± 0.09**** | 80.86 ± 10.29 | 10.04 ± 0.15* | -0.39 ± 0.03**** | 90.89 ± 12.79 | 7.57± 0.06**** | -1.00 ± 0.03**** | 91.69 ± 9.87 |
| **C267A and C273A** | 111.82± 7.47 | 8.70 ± 0.08**** | -1.69 ± 0.01**** | 52.83 ± 6.21**** | 8.99 ± 0.07**** | -1.34 ± 0.01**** | 58.43 ± 5.41**** | 6.83 ± 0.09**** | -1.98 ± 0.02**** | 41.35 ± 8.86**** |
| **P268A** | 94.55 ± 9.05 | 9.19 ± 0.07**** | -0.89 ± 0.07**** | 67.07 ± 6.10**** | 9.91± 0.10*** | -0.63 ± 0.01**** | 75.55 ± 8.53**** | 8.23 ± 0.14*** | -0.73 ± 0.10**** | 45.89 ± 15.69**** |
| **F277A** | 92.22 ± 6.61 | 9.83 ± 0.07* | -0.57 ± 0.07*** | 88.78 ± 3.24 | 10.45 ± 0.03 | -0.15 ± 0.03 | 89.42 ± 1.25 | 9.18 ±0.03 | -0.34 ± 0.04 | 101.33 ± 0.91 |
| **F280A** | 88.52 ± 3.75 | 8.81 ± 0.24**** | -1.31 ± 0.38**** | 108.36 ± 1.48 | 10.04 ± 0.16** | -0.26 ± 0.03*** | 107.48 ± 0.83 | 8.98 ± 0.22 | 0.44 ± 0.32** | 109.48 ± 1.29 |
| **L284A** | 100.28 ± 7.24 | 9.16 ± 0.08**** | -1.23 ± 0.16**** | 100.02 ± 3.74 | 10.63 ± 0.07 | -0.05 ± 0.03 | 100.49 ± 2.29 | 7.80 ± 0.08**** | -1.72 ± 0.05**** | 99.66 ± 4.72 |
| **D294A** | 161.95 ± 4.67**** | 8.32 ± 0.09**** | -2.08 ± 0.05**** | 58.09 ± 8.52**** | N.D. | N.D. | N.D. | N.D. | N.D. | N.D. |
| **D294N** | 196.64 ± 9.14**** | 10.48 ± 0.05 | 0.08 ± 0.05 | 106.32 ± 1.79 | N.D. | N.D. | N.D. | N.D. | N.D. | N.D. |
| **H301A** | 94.65 ± 8.54 | 10.51 ± 0.04 | 0.16 ± 0.06 | 96.88 ± 1.13 | N.D. | N.D. | N.D. | N.D. | N.D. | N.D. |
| **E304A and R307A** | 99.97 ± 3.96 | -9.51 ± 0.05**** | -0.83 ± 0.01**** | 72.76 ± 3.95*** | 9.75 ± 0.06**** | -0.70 ± 0.09**** | 76.43 ± 5.27**** | 7.96 ± 0.04**** | -0.91 ± 0.06**** | 27.89 ± 6.92**** |
| **ΔHelix 8** | 99.80 ± 10.12 | N.A. | N.A. | N.A. | N.A. | N.A. | N.A. | N.A. | N.A. | N.A. |

cAMP accumulation data were analyzed using a three-parameter logistic equation to determine $pEC_{50}$ and $E_{max}$ values. $pEC_{50}$ represents the negative logarithm of agonist concentration that produces half maximal response. $E_{max}$ is the maximal response as predicted by the dose-response curves. $E_{max}$ values for mutants were expressed as a percentage of the WT MC1R. Cell surface expression was assessed by FACS to detect the N-terminal Flag epitope label on the receptor and normalized to the WT receptor (shown as percentage). All data were expressed as means ± S.E.M. of at least three independent experiments. One-way ANOVA was used to determine statistical significance (*P< 0.05, **P< 0.01, ***P< 0.001 and ****P< 0.0001). N.D., not determined. N.A., not applicable.

**Table S6 Ligand binding affinities of WT and mutated MC1Rs.**

| MC1R | α-MSH | | Afamelanotide | | SHU9119 | |
|---|---|---|---|---|---|---|
| | $pK_i$ | $\Delta pK_i$ | $pK_i$ | $\Delta pK_i$ | $pK_i$ | $\Delta pK_i$ |
| **WT** | 8.47 ± 0.08 | 0 | 8.70 ± 0.06 | 0 | 8.64 ± 0.05 | 0 |
| **E94A** | 6.12 ± 0.95**** | -2.66 ± 0.31**** | N.D. | N.D. | N.D. | N.D. |
| **I98A** | 7.47 ± 0.15* | -1.00 ± 0.11*** | N.D. | N.D. | N.D. | N.D. |
| **D117A** | 6.27 ± 0.29**** | -2.22 ± 0.06**** | N.D. | N.D. | N.D. | N.D. |
| **D121A** | 6.47 ± 0.40**** | -2.01 ±0.14**** | N.D. | N.D. | N.D. | N.D. |
| **M128A** | 8.14 ± 0.10 | -0.34 ± 0.08 | N.D. | N.D. | N.D. | N.D. |
| **D184A-3×GSA-H185A** | 7.61 ± 0.17 | -0.69 ±0.05 | 8.69 ± 0.09 | 0.14 ± 0.08 | 8.52 ± 0.04 | -0.12 ± 0.06 |
| **L192A** | 8.13 ± 0.26 | -0.35 ± 0.10 | N.D. | N.D. | N.D. | N.D. |
| **H260A** | 7.58 ± 0.08 | -0.70 ± 0.04 | 9.17 ± 0.05 | 0.26 ± 0.28 | 9.29 ± 0.08 | 0.66 ± 0.13* |
| **C267A and C273A** | 8.38 ± 0.31 | 0.08 ± 0.12 | 8.52 ± 0.34 | -0.35 ± 0.09 | 8.84 ± 0.31 | 0.20 ±0.21 |
| **F280A** | 7.85 ± 0.04 | -0.66 ± 0.11* | N.D. | N.D. | N.D. | N.D. |

Competitive ligand binding results derived from three independent experiments performed in triplicate were normalized (nonspecific binding for 0% and total binding for 100%). pKi values were extracted from curve fitting and presented as means ± S.E.M. One-way ANOVA was used to determine statistical significance (*P< 0.05, **P< 0.01, ***P< 0.001, ****P< 0.0001). N.D., not determined.
